# High-fidelity approximation of grid- and shell-based sampling schemes from undersampled DSI using compressed sensing: Post mortem validation

**DOI:** 10.1101/2021.02.11.430672

**Authors:** Robert Jones, Chiara Maffei, Jean Augustinack, Bruce Fischl, Hui Wang, Berkin Bilgic, Anastasia Yendiki

## Abstract

While many useful microstructural indices, as well as orientation distribution functions, can be obtained from multi-shell dMRI data, there is growing interest in exploring the richer set of microstructural features that can be extracted from the full ensemble average propagator (EAP). The EAP can be readily computed from diffusion spectrum imaging (DSI) data, at the cost of a very lengthy acquisition. Compressed sensing (CS) has been used to make DSI more practical by reducing its acquisition time. CS applied to DSI (CS-DSI) attempts to reconstruct the EAP from significantly undersampled q-space data. We present a post mortem validation study where we evaluate the ability of CS-DSI to approximate not only fully sampled DSI but also multi-shell acquisitions with high fidelity. Human brain samples are imaged with high-resolution DSI at 9.4T and with polarization-sensitive optical coherence tomography (PSOCT). The latter provides direct measurements of axonal orientations at microscopic resolutions, allowing us to evaluate the mesoscopic orientation estimates obtained from diffusion MRI, in terms of their angular error and the presence of spurious peaks. We test two fast, dictionary-based, L2-regularized algorithms for CS-DSI reconstruction. We find that, for a CS acceleration factor of R=3, i.e., an acquisition with 171 gradient directions, one of these methods is able to achieve both low angular error and low number of spurious peaks. With a scan length similar to that of high angular resolution multi-shell acquisition schemes, this CS-DSI approach is able to approximate both fully sampled DSI and multi-shell data with high accuracy. Thus it is suitable for orientation reconstruction and microstructural modeling techniques that require either grid- or shell-based acquisitions. We find that the signal-to-noise ratio (SNR) of the training data used to construct the dictionary can have an impact on the accuracy of CS-DSI, but that there is substantial robustness to loss of SNR in the test data. Finally, we show that, as the CS acceleration factor increases beyond R=3, the accuracy of these reconstruction methods degrade, either in terms of the angular error, or in terms of the number of spurious peaks. Our results provide useful benchmarks for the future development of even more efficient q-space acceleration techniques.

## 1. Introduction

Diffusion magnetic resonance imaging (dMRI) has played an integral role in the study of human brain circuitry *in vivo* by enabling non-invasive investigation of tissue architecture (Le Bihan et al. 1986). The molecular displacements resulting from water diffusion can be estimated from dMRI measurements acquired with a pulsed gradient spin-echo (PGSE) sequence (Stejskal and Tanner 1965). Diffusion tensor imaging (DTI), the seminal approach for quantitative reconstruction of 3D water molecule displacement (Basser, Mattiello, and LeBihan 1994b, 1994a), assumes a 3D Gaussian displacement distribution and thus can only model a single fiber population in each voxel. One of the techniques that were introduced to resolve multiple intravoxel fiber populations is diffusion spectrum imaging (DSI), which relies on an acquisition that samples the entire q-space on a Cartesian grid (Wedeen et al. 2005). The diffusion ensemble average propagator (EAP), or the 3D probability density function (PDF) of spin displacements in a voxel, can be recovered directly from a Fourier transform (FT) of the Cartesian q-space signals, and yields a plethora of information describing the angular and radial features of diffusion (Wedeen et al. 2008; Hagmann et al. 2008).

We have previously used a DSI acquisition to collect data for our post mortem validation studies (Jones et al. 2020, Grisot et al. 2021). The benefit of this acquisition is that it yields a dataset that is densely sampled in q-space and can thus be used to generate data with other sampling schemes (such as single- or multi-shell) with q-space resampling. We have shown that this resampling approach can approximate q-shell data from data collected on a grid in q-space with high accuracy (Jones et al. 2020). Thus, from a single scan, we can generate data with multiple q-space sampling schemes and perform systematic, side-by-side comparisons, as we did most recently in the IronTract Challenge (Maffei et al. 2020, Maffei et al. 2021). Given that long acquisition times are one of the main challenges in the post mortem validation of dMRI (Yendiki et al. 2021), any opportunity to further reduce the scan time would be beneficial, as it would allow us to scan more samples or to scan the same samples at higher spatial resolution in the same amount of time.

Much recent work has been devoted to accelerating MR acquisitions. Some approaches modify imaging sequences to allow multiple image slices to be acquired simultaneously. Multi-slice parallel imaging techniques, such as simultaneous multi-slice (SMS) (Kawin Setsompop et al. 2018; Kawin Setsompop, Cohen-Adad, et al. 2012; Kawin Setsompop, Gagoski, et al. 2012), simultaneous image refocusing (SIR) (Reese et al. 2009), and multiplexed (SMS+SIR) echo planar imaging (EPI), have played a crucial role in reducing dMRI scan times down to reasonable lengths. However, they do not address the large number of q-space samples required by DSI. Compressed sensing (CS) has been applied to DSI (CS-DSI) to achieve q-space acceleration. CS theory exploits transform sparsity to recover signals from sub-Nyquist acquisitions (Donoho 2006; Lustig et al. 2008; Lustig, Donoho, and Pauly 2007). CS-DSI undersamples in q-space and reconstructs the missing samples with CS, allowing for a reduction in acquisition time directly proportional to the CS acceleration factor. Combining CS-DSI with SMS or multiplexed EPI can provide even higher accelerations (K. Setsompop et al. 2013), and render CS-DSI a practical diffusion protocol for large-scale *in vivo* population studies (Tobisch et al. 2018).

Several methods for CS-DSI reconstruction have been proposed. Menzel et al. (2011) used wavelet and total variation (TV) penalties on PDFs combined with random Gaussian undersampling patterns, and concluded that angular and radial diffusion properties were preserved at R=4 acceleration. Paquette et al. (2015) performed a joint comparison of different wavelet-based sparsifying transforms and q-space sampling strategies, and found the best results when using a “uniform-angular, random-radial” undersampling mask combined with discrete wavelet transform (DWT). Both Menzel et al. and Paquette et al. used fixed transforms to generate sparse signal representations. Conversely, Bilgic et al. (2012) proposed CS-DSI using adaptive PDF dictionaries, combining the K-SVD algorithm (Aharon, Elad, and Bruckstein 2006) for dictionary training with the FOcal Underdetermined System Solver (FOCUSS) algorithm (Gorodnitsky and Rao 1997) to solve the CS problem. While this approach yielded reduced reconstruction errors compared to fixed transforms, the iterative FOCUSS reconstruction resulted in full brain computation times on the order of days. This bottleneck was addressed in subsequent work (Bilgic et al. 2013), where two dictionary-based, L2-regularized methods were introduced that reduce computation times down to seconds per slice. They provide fast, simple formulations while preserving reconstruction quality compared to Dictionary-FOCUSS. One of the key findings from (Bilgic et al. 2013) was that forcing the PDFs to remain in the range of a dictionary was more important than the sparsity constraints imposed on the transform coefficients. In other words, the key to good reconstructions lies in the prior information encoded in a dictionary, and not the regularization norm that is applied on the dictionary transform coefficients. We note that the theoretical foundation of CS concerns signal recovery using the L1-norm, and that the aforementioned CS-DSI methods use the L2-norm. With this in mind, we will use CS-DSI to refer to the umbrella of algorithms that reconstruct DSI from undersampled q-space.

Dictionary-based CS-DSI is a promising approach, but nonetheless has aspects that require further investigation. One of these areas is the effect of dMRI signal-to-noise ratio (SNR) on the CS algorithms and dictionary learning, which is yet to be well characterized (Bilgic et al. 2012). Determining how the SNR of the training data influences reconstructions and the minimum SNR level required for high-quality CS reconstruction would provide critical insights for the development of CS-DSI protocols. Previously, CS-DSI reconstruction fidelity was assessed *in vivo*. In that case, the only available ground truth is a the fully sampled DSI dataset from the same brain (Bilgic et al. 2013; Bilgic et al. 2012; Menzel et al. 2011). A drawback of this approach is that the fully sampled DSI data are inherently corrupted with noise, particularly at high b-values. Bilgic et al. (2013) ameliorated this issue by sampling a few q-space locations 10 times, and using these low-noise references to evaluate reconstructions. Interestingly, they found that CS reconstructions exhibited lower errors than the fully sampled (1-average) data when compared to the 10-average data. These results exemplify potential denoising benefits of CS-DSI with respect to fully sampled data collected at the same SNR, but also highlight inadequacies of taking the fully sampled data as ground-truth. Ultimately, our goal is to use the data to characterize the underlying white-matter fiber geometries. Thus, it is important to know how great an error with respect to the fully sampled data we can tolerate in our CS reconstructions, without compromising accuracy with respect to the true fiber orientations. For this purpose, we need independent measurements of fiber orientations from a modality that does not rely on water diffusion.

Here, we address these questions with a validation study of CS-DSI reconstructions in *ex vivo* human brain. Our goal is two-fold. First, we perform the first validation study that assesses the accuracy of CS-DSI with respect to ground-truth measurements of axonal orientations from optical imaging. This is of relevance to the use of CS-DSI in large-scale population imaging to study white-matter organization across the human lifespan (Tobisch et al. 2018). Second, we investigate the ability of CS-DSI reconstruction to approximate multiple q-space sampling schemes, including grid- and shell-based. This is of relevance to both in vivo population studies and ex vivo validation studies, where the flexibility to use orientation reconstruction methods and microstructural models that require either type of sampling scheme is an asset.

Specifically, we image human white-matter samples with high-resolution DSI at 9.4T and with polarization-sensitive optical coherence tomography (PSOCT) (De Boer et al. 1997). The latter is a technique in the family of label-free 3D optical imaging methods, which provide direct measurements of fiber orientations at microscopic resolution (Wang et al. 2018; Wang et al. 2011) and have emerged in recent years as a powerful tool for the validation of the mesoscopic-resolution diffusion orientations obtained from dMRI (see Yendiki et al., (2021) for a review). We have recently demonstrated the ability of PSOCT to resolve complex fiber configurations, including interdigitated crossing fibers, distinct (non-interdigitated) crossing fibers, and branching fibers, in human brain tissue (Jones et al. 2020). Numerous other studies have employed PSOCT to resolve crossing fibers in biological tissues (Villiger et al. 2018; Ruiz-Lopera et al. 2021; Yao and Duan 2020; Guo et al. 2004; Fan and Yao 2013), including the mouse brain (Lefebvre et al. 2021) and human brain (Boas et al. 2017; Wang et al. 2018).

We undersample the DSI data and use the dictionary-based techniques from Bilgic et al. (2013) to perform CS reconstructions, with dictionaries trained on DSI data from three different *ex vivo* human brain blocks to determine generalizability. We investigate if fiber orientations estimated from CS reconstructed data can achieve the same accuracy as those estimated from the fully sampled data, by comparing both to the ground-truth fiber orientations measured by PSOCT. We examine the influence of the SNR of the training or test data on the accuracy of the diffusion orientation estimates. We show that, for an acceleration factor of R=3, which corresponds to 171 diffusion-encoding gradient directions, the accuracy of CS-DSI is very similar to that of fully sampled DSI. Our findings indicate that, with an acquisition time similar or shorter than typical high angular resolution multi-shell scans, CS-DSI data can be used to approximate both fully sampled DSI and multi-shell data with high accuracy. This provides the flexibility to take advantage of the high angular accuracy of DSI (Daducci et al. 2013; Jones et al. 2020; Maffei et al. 2020, 2021) while also allowing microstructural analyses either on q-shells or on the full EAP. For *in vivo* population studies, this gives access to a wider range of potential biomarkers. For *ex vivo* validation studies, it facilitates the comparison of multiple acquisition schemes from a single scan. Finally, we find that, when the acceleration factor of CS-DSI is increased beyond R=3, performance deteriorates, increasing either the angular error or the number of spurious peaks. These results provide a benchmark for the future development of more efficient q-space acceleration techniques.

## 2. Methods

### 2.1. Sample identification

The samples used in this study were extracted from two human brain hemispheres that were obtained from the Massachusetts General Hospital Autopsy Suite and fixed in 10% formalin for at least two months. Demographic information about the hemispheres is given in Table 1. Three samples were extracted from different anatomical locations of the hemispheres. Sample 1A (Figure 1A) was cut from brain 1 and was approximately 3×2×2 cm. It contained an area of deep white matter (WM) that included the corpus callosum (CC), the internal and external capsules (IC and EC, respectively), the caudate nucleus and the putamen. Sample 1B (Figure 1B) was taken from a different region of brain 1. The block was approximately 3×2×2 cm and contained an area of deep WM including the posterior internal capsule, putamen, and thalamus. Sample 2 (Figure 1C) was cut from brain 2 and was sized approximately 2×2×3 cm. The superior part of the block contained the anterior part of the superior frontal gyrus, the medial side included the cingulate sulcus, and the lateral side contained parts of the corticospinal tract and dorsal superior longitudinal fasciculus (SLF-I).

**Table 1.** Demographic information on post mortem human hemispheres.

| Brain | Age | Gender | Laterality | Cause of death | Diagnosis | PMI |
| --- | --- | --- | --- | --- | --- | --- |
| 1 (samples A and B) | 43 | F | Right | Cardiac arrest | Cognitive control | <24 h |
| 2 | 70 | M | Left | Coronary artery disease and multiorgan failure | Severe coronary artery disease, myocardial infarction, hypertension, hyperlipidemia, mild hypoxic/ischemic changes | 24 h |

**Figure 1.**
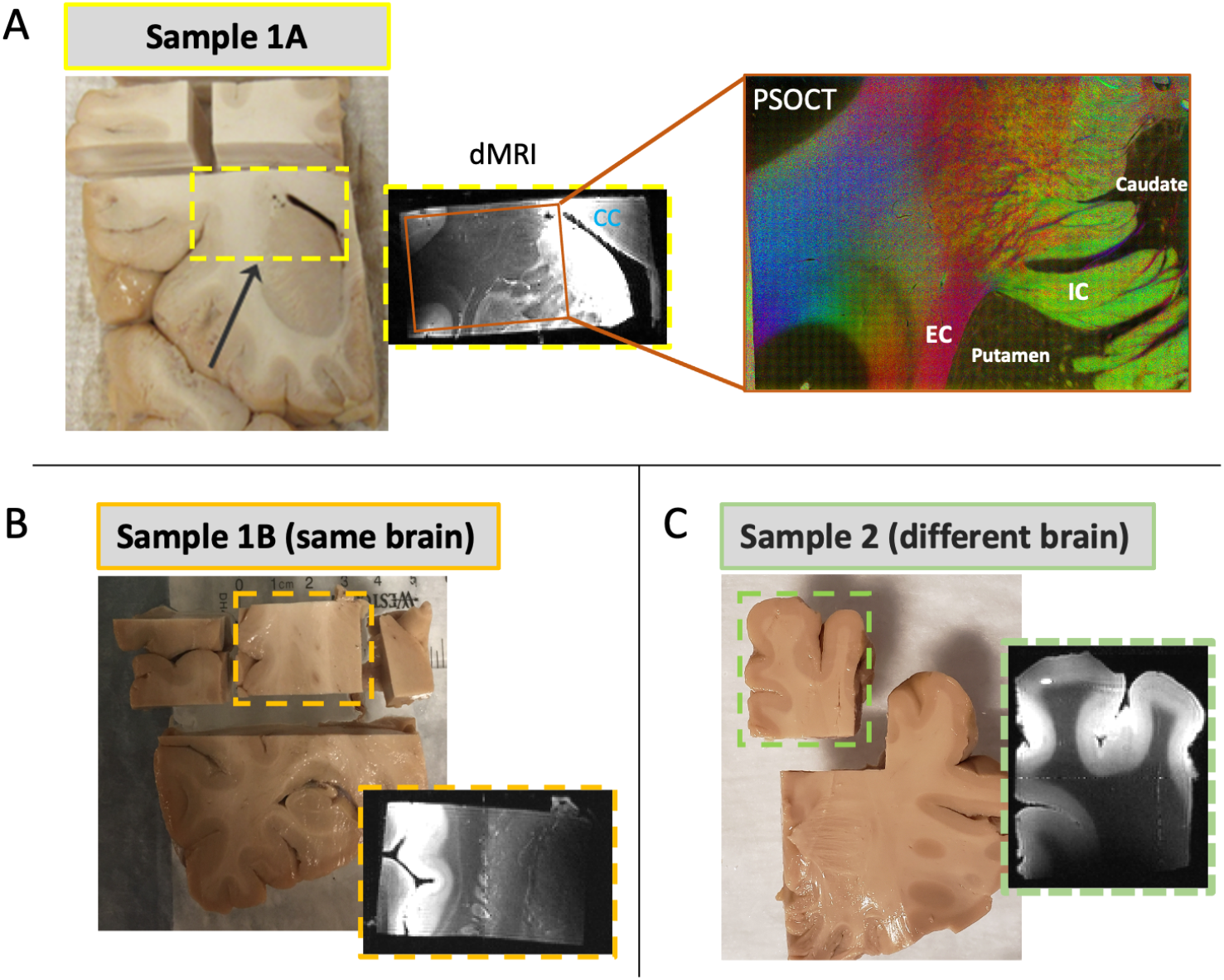
Overview of sample identification and data acquisition. *Ex vivo* human brain samples were extracted from coronal slabs (dashed rectangles) and dMRI data were acquired at 9.4T. Single slices from b=0 scans are shown for each sample. Samples 1A and 1B were cut from different anatomical locations of the same brain, and sample 2 was extracted from a different brain. Following dMRI, a piece of sample 1A was cut and imaged with PSOCT (A, right). Sample 1A, which contained the corpus callosum (CC), internal capsule (IC), external capsule (EC), caudate and putamen, was used as the test dataset. Each of samples 1A, 1B, and 2 were used as training datasets.

### 2.2. Diffusion MRI

#### 2.2.1. Data acquisition

All three *ex vivo* samples were scanned in a small-bore 9.4T Bruker Biospec system with gradients capable of G_max_=480 mT/m. Prior to scanning, each block was placed in a plastic syringe filled with Fomblin (perfluoropolyether) and all air was removed. The dMRI data were acquired using a spin echo 3D single-shot EPI sequence with G_max_=393 mT/m, TR=750 ms, TE=43 ms, GRAPPA factor 2, matrix size 136×136×176, and 250 μm isotropic resolution. We used a DSI sampling scheme consisting of one b=0 image and 514 gradient directions arranged on a Cartesian lattice in q-space and zero padded to an 11×11×11 grid (Wedeen et al. 2005). Diffusion encoding was applied with b_max_=40000 s/mm^2^, δ=15 ms, and Δ=21 ms, corresponding to q_max_=250 mm^−1^. The total acquisition time was approximately 48 hours. We will refer to the datasets containing all 515 diffusion volumes as the “fully sampled” or “FS” data.

A 4-channel phased array surface receive (Rx) coil was used in dMRI acquisitions (diagramed in Figure 2A, bottom), leading to a decrease in coil sensitivity and SNR as the distance between the sample and Rx coil increased. The coronal planes of the dMRI data were approximately parallel to the surface coil, so coronal slices closer to the coil had higher signal than the slices farther away. Figure 2A (top) shows three example coronal slices from the b=0 volume of sample 1A, each with varying distances from the surface Rx coil. To quantify the dMRI SNR, we used the b=0 signal intensities from a region of interest (ROI) in the deep WM. The SNR of each coronal slice was calculated as the mean signal (Figure 2B, solid black line) divided by the standard deviation of the signal intensities from the ROI (Figure 2B, solid gray line). To relate the SNR across slices to the sensitivity profile of the coil, we fit a linear model to the calculated SNR values to obtain an expression for SNR as a function of slice (Figure 2B, dotted green line). For the dMRI slice numbering of each sample, we will refer to slice 1 as the slice closest to the surface coil, with increasing slice numbers indicating increased distance from the coil.

**Figure 2.**
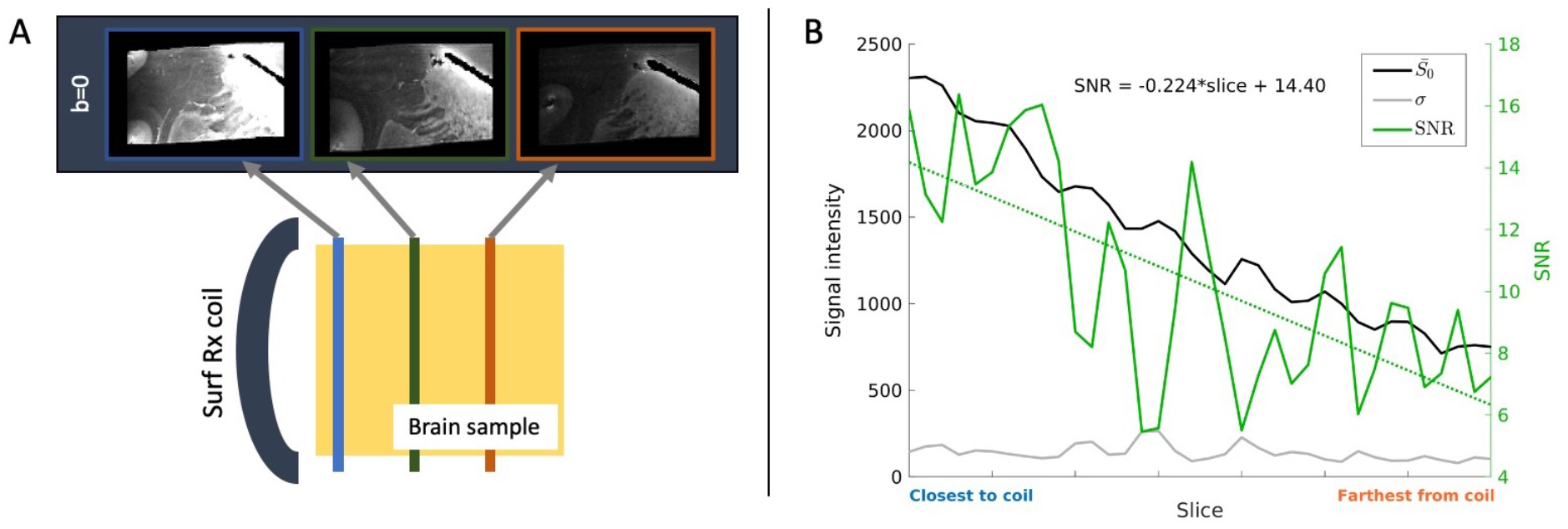
dMRI acquisition and SNR. **(A)** Bottom: A surface receive coil was used in dMRI acquisitions. Top: This led to a decrease in sensitivity with increasing distance from the coil, as shown in coronal slices from the sample 1A b=0 volume. The same intensity scaling was used for all three slices. **(B)** The b=0 SNR in each coronal slice of sample 1A (solid green line) was calculated from the mean (black line) and standard deviation (gray line) of signal intensities from a deep WM ROI. A linear regression was performed to find the slope of SNR as a function of slice (dotted green line). The left y-axis shows the signal and noise intensities, and the right y-axis shows SNR values. The x-axis in indexed by slice number.

#### 2.2.2. Compressed sensing reconstruction

CS reconstructions were performed using two dictionary-based CS-DSI methods previously introduced by Bilgic et al. (2013). One is PCA-based reconstruction (PCA), and the other is Tikhonov-regularized pseudoinverse reconstruction using the training set of PDFs as the dictionary (PINV). CS undersampling masks were generated for acceleration factors R=3, 5, and 9 using a variable-density power-law function (Lustig, Donoho, and Pauly 2007; Bilgic et al. 2013). Nine CS masks with different sampling patterns were created for each acceleration factor. Retrospective undersampling was applied to fully sampled q-space data, followed by CS reconstruction using either PCA or PINV. Each reconstruction method had one free parameter: the number of principal components *T* and the Tikhonov regularization parameter *λ*, for PCA and PINV respectively. The optimal parameter for each dictionary was determined using the parameter sweeping approach from Bilgic et al. (2013), in which the training data were undersampled and reconstructed using a range of parameters, and the one yielding the lowest root-mean-square error (RMSE) in PDFs (described in section 2.4.2.) when compared to the fully sampled data was selected. This procedure was performed separately for each combination of dictionary and acceleration factor.

Reconstructions and analyses were performed in MATLAB (R2019a, 9.6) based on publicly available code (https://www.martinos.org/~berkin/software.html). Both PCA and PINV methods used a single matrix multiplication to reconstruct an entire slice, with computation times of ~10-15 seconds per slice (~6000 voxels per slice) on a workstation with a 3.4GHz Intel i7 processor, 8 cores, 32GB RAM. For full volume reconstructions (~50 slices), we compiled the MATLAB function into a standalone application and processed individual slices in parallel on a high-performance compute cluster. Simultaneously running each slice as a separate process with 8GB of allocated memory resulted in full volume reconstruction times between 2-10 minutes (depending on the number of available CPUs on the cluster).

#### 2.2.3. Fiber orientation reconstruction

At each dMRI voxel, we computed the orientation distribution function (ODF), *i.e.*, the marginal PDF of the diffusion orientation angle. Diffusion tractography algorithms use the ODFs, rather than the full PDFs, to reconstruct WM bundles. In DSI, ODFs are typically obtained by interpolating the PDFs onto uniform radial vertices and summing them along each radial projection. The truncation of q-space causes ringing in the PDFs, which introduces artifacts into the ODFs. One approach to mitigating these artifacts is to apply a windowing function on the q-space data, like a Hanning filter (Wedeen et al. 2005). This smooths the signal decay at the edges of q-space but diminishes the contributions of high-frequency diffusion terms, potentially oversmoothing the PDFs and ODFs and reducing angular resolution. An alternative approach is to use unfiltered q-space data and carefully define the starting and ending displacement distances for integration of the PDF (Tian et al. 2016; Paquette, Gilbert, and Descoteaux 2016; Lacerda et al. 2016). This way, one can restrict the integration range so that PDF ringing is omitted from ODF computations without having to taper the high frequency q-space data. We used the latter approach for ODF reconstructions.

DSI ODF reconstructions were performed in Python with the *Dipy* (version 1.3.0.) library (Garyfallidis et al. 2014). For each voxel, the q-space data were zero-padded to a 75×75×75 grid and the 3D FFT was applied to obtain the PDF, with negative PDF values clipped to zero. For ODF reconstruction, we used a PDF integration lower bound of *r*_*start*_ = 10 and upper bound of *r*_*end*_ = 16 (which were roughly 0.27x and 0.43xFOV), a radial step size of *r*_*step*_ = 0.05, and a radial weighting factor of 2. ODF peaks were extracted using a maximum of 3 peaks per voxel, minimum peak separation angle of 25° and minimum peak magnitude of 0.05. The same parameters were used for all DSI ODF reconstructions.

We also fit the DTI model to the fully sampled data using the FSL command *dtifit* and extracted the orientations of the primary eigenvectors of the tensors. These served as a baseline for “worst-case” dMRI orientation accuracy, *i.e.*, where only a single fiber population can be resolved in each voxel. We will refer to the fully sampled DTI and DSI orientations as FS-DTI and FS-DSI, respectively.

#### 2.2.4. Resampling onto q-shells

Next, we investigated how well we could approximate q-shell data from CS-DSI acquisitions by comparing data that were acquired with CS-DSI scheme and resampled onto q-shells to data from the same sample that were acquired on q-shells. Following CS reconstruction, the CS-DSI q-space data were resampled onto shells in q-space using the approach described in (Jones et al. 2020). Briefly, the q-space samples arranged on shells were approximated by using a nonuniform fast Fourier transform (NUFFT) with min-max interpolation (Fessler and Sutton 2003) to interpolate the DSI q-space samples arranged on a dense grid. Multi-shell data from sample 1B that were generated by resampling FS-DSI [NUFFT(FS)] and CS-DSI [NUFFT(CS)] q-space data were compared to multi-shell q-space data of sample 1B that were acquired during a separate scan.

The multi-shell acquisition of sample 1B used identical imaging parameters as the FS-DSI acquisition (described in section 2.2.1.), except that it consisted of one b=0 image and 256 directions arranged on 3 q-shells, with 64, 64, 128 directions and b=4,000, 12,000, 20,000 s/mm^2^, respectively. FS-DSI and CS-DSI q-space data from sample 1B were resampled onto the same 3 shells and compared to the acquired multi-shell data. The CS-DSI reconstruction tested here used the PCA method at acceleration R=3 and training data from sample 2.

As previously (Jones et al. 2020), resampling fidelity was assessed using the relative normalized root mean square error (rNRMSE) between the acquired and resampled q-space data. This accounts for the fact that the acquired and resampled data come from different scan sessions, and thus some differences between them are expected as they contain different noise realizations. This is done by normalizing the NRMSE of the q-space images by the NRMSE of the b=0 images from each acquisition. Thus the rNRMSE quantifies the extent to which resampling introduces additional error, with respect to what the error would have been due to noise alone. For each diffusion-weighted volume, we compared the average rNRMSE across all WM voxels for the resampled FS-DSI data [NUFFT(FS-DSI)] and resampled CS-DSI data [NUFFT(CS-DSI)].

We also investigated the effect of q-space resampling on indices extracted from microstructural dMRI models. For each of the three multi-shell datasets, i.e., acquired, resampled NUFFT(FS), and resampled NUFFT(CS), the diffusion kurtosis imaging (DKI) model was fit to the first two shells (b=4,000 and 12,000 s/mm^2^) using an ordinary least-squares approach implemented in *Dipy* (version 1.3.0). Microstructural indices were extracted from the fitted tensors, namely the fractional anisotropy (FA) of the diffusion tensor, the orientation of the primary eigenvector of the diffusion tensor, the mean kurtosis (MK), the radial kurtosis (RK) and the axial kurtosis (AK).

### 2.3. PSOCT

#### 2.3.1. Data acquisition

Following dMRI acquisition of sample 1A, a piece of the tissue block was extracted for imaging with PSOCT (Figure 1A, right). For a thorough review of PSOCT and its applications, see de Boer et al. (2017). Briefly, PSOCT uses polarized light to probe tissue birefringence and obtain undistorted, direct measurements of in-plane fiber orientations at microscopic resolutions. Thus PSOCT has been used as a reference modality for the validation of dMRI-derived fiber orientation estimates (Wang et al. 2014; Jones et al. 2020). As these studies have shown, PSOCT can resolve complex fiber patterns at a scale that is not accessible with dMRI. That is because the spatial resolution of PSOCT is 2-3 orders of magnitude higher than that of dMRI, and thus much closer to the caliber of axons (<1-10 μm; (Liewald et al. 2014)).

Details on our setup, acquisition, and analysis for PSOCT were previously described (Jones et al. 2020). Briefly, the sample was imaged with a polarization maintaining fiber (PMF) based, spectral domain PSOCT system developed in-house (Wang et al. 2016). The light source consisted of two broadband super-luminescent diodes (Thorlabs Inc., LSC2000C), with a center wavelength of 1300 nm and a bandwidth of 170 nm. The axial resolution was 2.9 μm in tissue and the lateral resolution was estimated at 3.5 μm. PSOCT produces measurements of optic axis orientation, which represent the in-plane (2D) orientation of the fiber axis. We downsampled the PSOCT data to an in-plane resolution of 10 μm to facilitate data processing. To cover the entire tissue surface, 1120 tiles (FOV = 1 mm^2^) were acquired using a snaked configuration scheme with 50% overlap between adjacent tiles. The tiles were stitched using the Fiji software (Schindelin et al. 2012; Preibisch, Saalfeld, and Tomancak 2009). A vibratome cut off a 75 μm slice after the superficial region of the tissue block was imaged, which consequently allowed deeper regions to be exposed by PSOCT. There were 63 total slices acquired for the sample block. One critical advantage of the technique is that PSOCT images the blockface of the tissue before slicing (Wang et al. 2018). This avoids the nonlinear slice distortions that are present in traditional histological techniques, where slices are imaged after they are cut. As a result, we can simply stack the slices into a volume.

#### 2.3.2. Cross-modal registration

The registration of PSOCT and dMRI images was performed as described in Jones et al. (2020). There, we examined whether our dMRI scans had nonlinear distortions that might affect image registration. We registered the dMRI b=0 volume to a structural MRI scan from the same sample, using either affine or nonlinear registration. We found that b=0 voxel intensities after affine vs. nonlinear registration were highly correlated (***ρ*** = 0.96, p < 0.001), indicating that nonlinear distortions were negligible and therefore affine alignment was sufficient.

Affine registration was used to align the dMRI fractional anisotropy (FA) and PSOCT retardance volumes. Retardance represents the phase delay between orthogonal polarization channels that is induced by birefringence, a property of anisotropic structures. The myelinated axons that compose WM bundles possess birefringent properties and are highlighted in the retardance, providing a tissue contrast similar to FA. We used a robust, inverse consistent registration method that detects and down-weighs outlier regions in the images (Reuter, Rosas, and Fischl 2010). Previously, Wang et al. (2014) reported a Dice coefficient of 0.96 between co-registered FA and retardance volumes with this approach, signifying the exceptional alignment that can be attained between the two contrasts.

We transformed the dMRI volumes to PSOCT space using nearest-neighbor interpolation, and also rotated the dMRI orientation vectors (see section 2.2.3 above) accordingly using the rotational component of the affine transformation. Finally, we projected the 3D dMRI vectors onto the 2D PSOCT imaging plane for the purposes of comparison to PSOCT optic axis measurements, which represent in-plane (2D) fiber orientations.

### 2.4. Error metrics

#### 2.4.1. Angular error with respect to PSOCT

We performed a voxel-wise comparison of dMRI and PSOCT orientations using absolute angular error as the accuracy metric, as in (Jones et al. 2020). In each voxel, we selected the peak of the diffusion ODF with the (2D) in-plane orientation that matched the corresponding PSOCT orientation most closely and computed the angular error between that peak and the PSOCT measurement. Thus, for a voxel with *N* dMRI ODF peaks, the absolute in-plane angular error *AE* was calculated as:

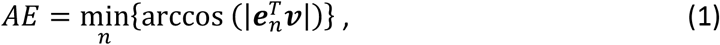

where ***e**_n_* is the unit vector along the *n^th^* dMRI peak orientation, and ***ν*** is the unit vector along the measured PSOCT orientation. Since the analysis was performed in 2D, the angular error could take values between 0° and 90°.

A WM mask was created to exclude all voxels where the retardance intensity was below 50% of the maximum retardance, and only voxels in this WM mask were considered when computing angular error metrics. PSOCT optic axis measurements rely on the birefringence of anisotropic processes, such as axon bundles in WM, but are not necessarily accurate in gray matter (GM) or voxels where fibers primarily project through the imaging plane, where the lack of apparent birefringence results in low retardance.

The PSOCT imaging plane was nearly parallel to the dMRI coronal plane, which itself was nearly parallel to the surface Rx coil. We exploited this arrangement, and the decreasing SNR of dMRI slices at increasing distance from the coil, to assess the accuracy of CS-DSI as a function of SNR.

#### 2.4.2. RMSE in PDFs with respect to FS-DSI

As a more conventional CS-DSI error metric, we also computed the normalized RMSE of the PDFs obtained from CS reconstruction with respect to those obtained from the fully sampled DSI data:

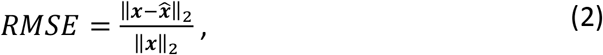

where ***x*** is the PDF from the fully sampled data, 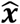 is the PDF from the CS reconstructed data, and ‖·‖_2_ is the L2-norm.

#### 2.4.3. Number of fiber populations per voxel

To gauge the specificity of dMRI fiber orientation estimates, we investigated the number of fiber orientations per voxel extracted from dMRI ODFs and from PSOCT measurements. For each dMRI reconstruction, we calculated the average number of detected ODF peaks (max of 3 in each voxel) across all WM voxels. To estimate the number of PSOCT fibers per voxel, we used an approach similar to the one described in Jones et al. (2020). First, PSOCT fiber orientation distributions (FODs) were constructed by generating histograms of the orientation measurements from all 0.01 mm PSOCT voxels within each 0.25 mm dMRI voxel, using a bin width of 5°. Then, the local maxima of the PSOCT FODs that had a height of at least 5% of the maximum bin and were separated by at least 10° were extracted using the *findpeaks* function in MATLAB. For consistency with dMRI, we also imposed a maximum of 3 PSOCT FOD peaks per voxel. The average number of PSOCT fibers per voxel was then calculated by averaging the number of PSOCT FOD peaks across all WM voxels in the volume.

### 2.5. CS-DSI validation experiments

A total of six PDF dictionaries were constructed, each trained on a single slice of fully sampled DSI data from sample 1A, sample 1B, or sample 2. For each sample we created two dictionaries, one from a high-SNR slice (slice 3) and one from a low-SNR slice (slice 13). We applied CS reconstruction to DSI data from sample 1A, undersampled by a factor of R=3, 5, or 9, using one of these dictionaries. We then computed DSI ODFs and extracted the orientations of the ODF peaks. We transformed the diffusion orientation vectors to PSOCT space, projected them onto the PSOCT plane, and calculated the absolute angular error with respect to PSOCT at each WM voxel.

Angular non-uniformities in the CS undersampling patterns may introduce directional biases into the ODFs, and thus affect our angular error computations. We accounted for this potential source of variability in our error metrics by repeating the CS reconstructions with 9 different CS undersampling masks, for each combination of dictionary and acceleration factor. Visualizations of the 3D angular distributions for all CS undersampling masks used in this study are provided in the supplementary information (Figures S1–S3).

#### 2.5.1. Effect of CS acceleration factor and training sample on angular error

We assessed the efficacy of the CS algorithms at different acceleration factors, by comparing CS reconstructions of data from sample 1A that had been undersampled by a factor of R=3, 5, and 9. In this comparison, we used the PCA and PINV methods with dictionaries trained on a high-SNR slice from each sample. The accuracy of dMRI orientations was quantified by the mean angular error across each PSOCT slice, as well as across all WM voxels in the PSOCT volume. This error was averaged over the reconstructions that were obtained with the 9 different CS undersampling masks. The mean angular error of FS-DSI was used as a reference for evaluating the quality of CS reconstructions.

#### 2.5.2. Effect of SNR on reconstruction error metrics

One goal of this study was to determine the influence of SNR on metrics of CS reconstruction quality. To this end, we calculated the b=0 SNR (as described in section 2.2.1.) for each dMRI slice and computed the average RMSE in PDFs (with respect to the fully sampled data) and the average angular error (with respect to PSOCT) across all WM voxels in each slice. Then, for each CS reconstruction, we performed a linear regression of RMSE or angular error against SNR. This was done for FS-DTI, FS-DSI, and CS-DSI. For CS-DSI, error metrics from each combination of acceleration, training sample, and method were calculated, each time averaging the mean errors over CS reconstructions from the 9 different undersampling masks.

#### 2.5.3. Effect of SNR on dictionary training

To evaluate the effect that the SNR of the training data has on CS reconstructions, voxels in sample 1A were reconstructed at acceleration R=3 with dictionaries trained on low-SNR data (slice 13) from each sample using PCA and PINV methods. These results were then compared to the corresponding results from the experiment described in section 2.5.1. that used high-SNR training data. The accuracy of dMRI orientations was quantified by the mean angular error across all WM voxels in the PSOCT volume. The CS reconstructions included here were performed with one randomly selected CS undersampling mask.

## 3. Results

### 3.1. Visual inspection

Figure 3 provides visualizations of dMRI and PSOCT orientations from a representative slice of sample 1A. Figure 3A shows fiber orientation maps as color-coded RGB images, where the color wheel shows the correspondence between pixel color and in-plane orientation. For FS-DSI and for CS-DSI at acceleration factors R=3, 5, and 9, the color maps show the orientations of the ODF peaks that most closely matched the PSOCT orientations in the same voxel. All CS-DSI results are shown for the same CS undersampling mask. For fully sampled DTI, the color maps show the orientations of the primary eigenvector of the diffusion tensor. Figure 3B shows heat maps of the absolute angular error between dMRI and PSOCT orientations in each voxel.

**Figure 3.**
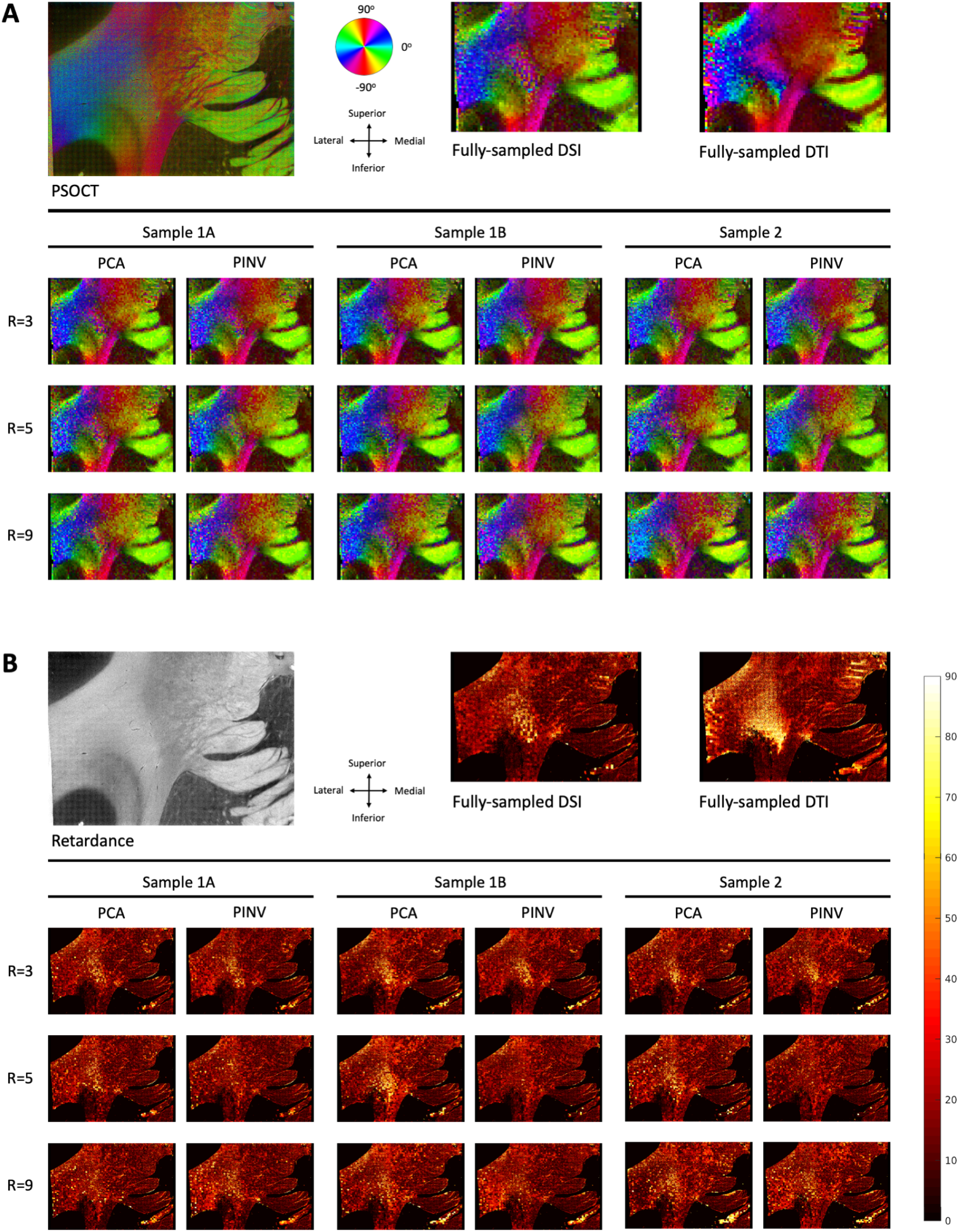
Fiber orientations estimated from dMRI vs. PSOCT. **(A)** Color-coded maps of PSOCT (top left) and dMRI orientations from fully sampled data (top right) and CS-DSI data (bottom). **(B)** Absolute angular error of dMRI orientations with respect to PSOCT. A WM mask was created by thresholding the PSOCT retardance (top left). The heat maps were masked to include only voxels classified as WM. CS-DSI reconstructions are shown for high-SNR training data and one of the CS undersampling masks.

Despite the large disparity in voxel size, there was good overall agreement between dMRI and PSOCT fiber orientation maps (Figure 3A). The dMRI maps showed the closest resemblance to PSOCT in the medial half of the slice, with greater differences in the lateral half of the slice. Examination of the angular error maps (Figure 3B) confirms that the greatest angular errors occurred in the middle and lateral regions of the slice. This distribution of errors was most obvious in the FS-DTI error map (Figure 3B, top right). The fully sampled (Figure 3B, top middle) and CS-DSI maps (Figure 3B, bottom) show similarly good agreement with PSOCT.

### 3.2. Effect of CS acceleration factor and training sample on angular error

Figure 4 shows bar plots of the mean angular errors of dMRI with respect to PSOCT, averaged over all WM voxels analyzed, from CS reconstructions of sample 1A with high-SNR training data from different samples and at different acceleration factors. Error bars show the standard error over different CS undersampling masks. The corresponding statistics are given in Table 2. The mean angular error of FS-DSI (green bar and dotted line) is shown as the benchmark for assessing CS-DSI results. The mean angular error of FS-DTI (red bar and dotted line) is shown as a worst-case scenario.

**Table 2.**
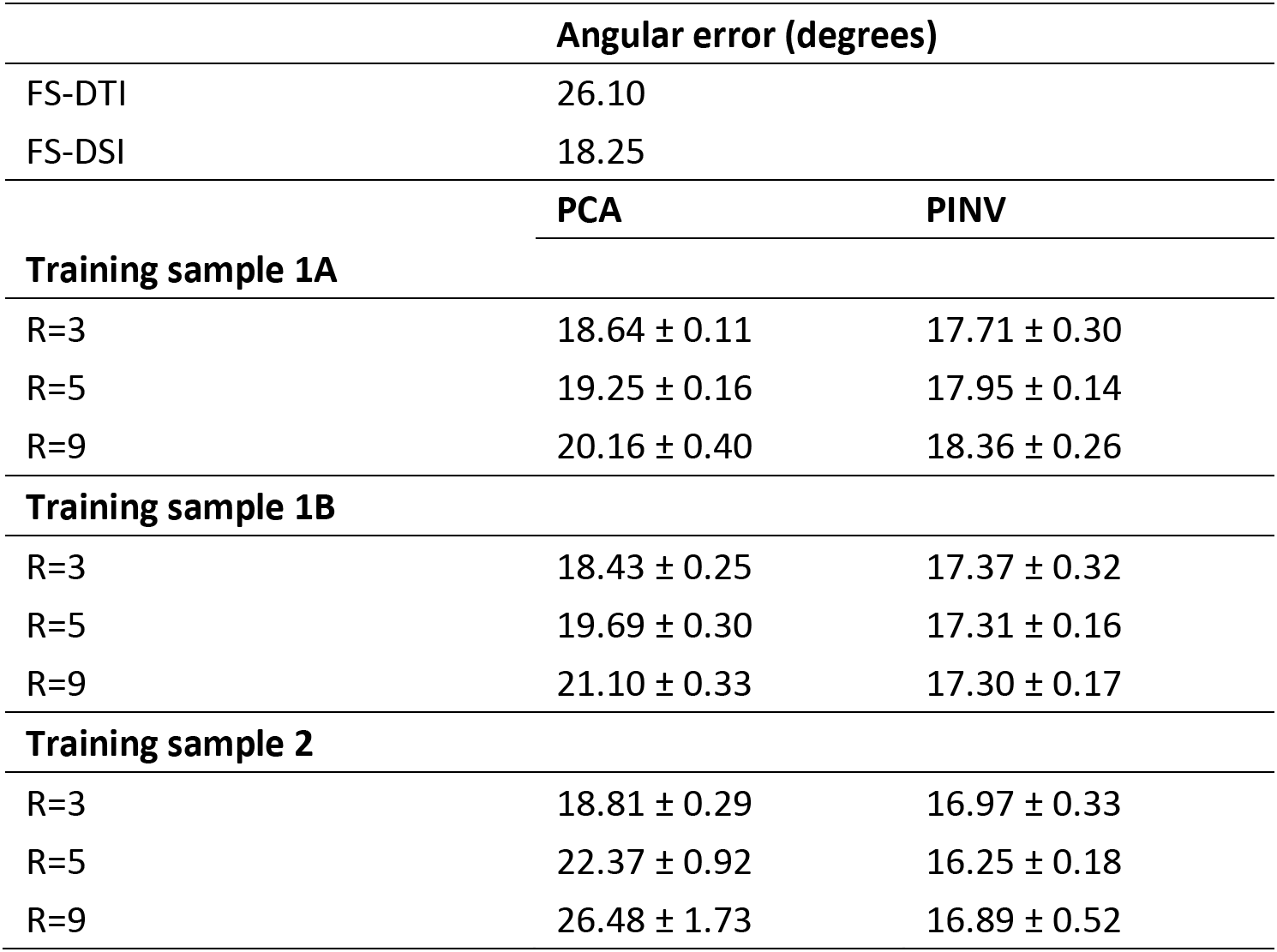
Angular error of dMRI orientations. Mean angular errors of dMRI with respect to PSOCT across all analyzed WM voxels. CS-DSI reconstructions used high-SNR training data from each sample. For CS-DSI, standard errors of the mean are also shown, computed over the 9 CS undersampling masks.

**Figure 4.**
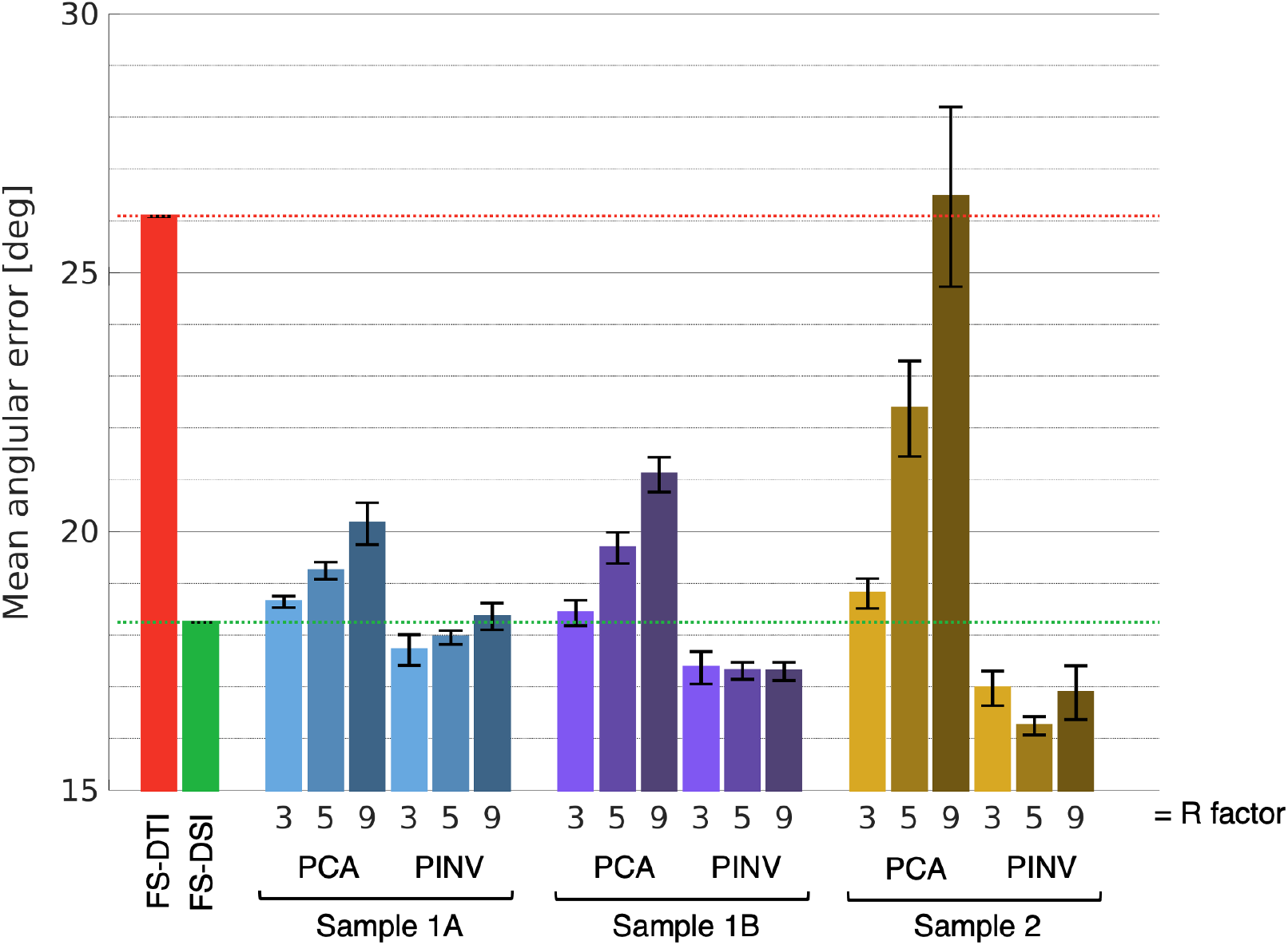
Mean angular error of dMRI with respect to PSOCT. The plots show average error over all WM voxels in sample 1A. For CS-DSI, the error was also averaged over 9 CS undersampling masks, with error bars representing standard error of the mean across these 9 masks. Results are grouped by training sample (sample 1A, blue; sample 1B, purple; sample 2, yellow) and CS method. Bar shades correspond to acceleration factor (3, 5, 9). All CS-DSI reconstructions used high-SNR training data from each sample. Mean angular errors from FS-DTI (red) and FS-DSI (green) are shown on the far right for comparison.

At an acceleration factor of R=3, CS-DSI achieved very similar angular error to FS-DSI (within +/−1.27 °), for both PCA and PINV reconstruction methods, and regardless of whether the training data came from the same or a different sample than the test data. For PCA reconstruction, the angular error increased with the acceleration factor. This increase was most dramatic in the (more realistic) scenario where the training data came from a different brain than the test data. Conversely, higher acceleration factors imparted only minor changes on the accuracy of PINV reconstructions.

We delved deeper into this difference between the performance of PCA and PINV by comparing the average number of ODF peaks per voxel from the data reconstructed by each method. The bar plot in Figure 5A (left) shows the average number of reconstructed ODF peaks per WM voxel from each reconstruction (for a maximum of 3 peaks per voxel), with error bars showing the standard error of the mean. CS-DSI results were averaged over the 9 undersampling masks used for CS reconstructions. The number of peaks per voxel from FS-DSI ODFs (green bar) and from PSOCT FODs (dashed red line) are shown for reference.

**Figure 5.**
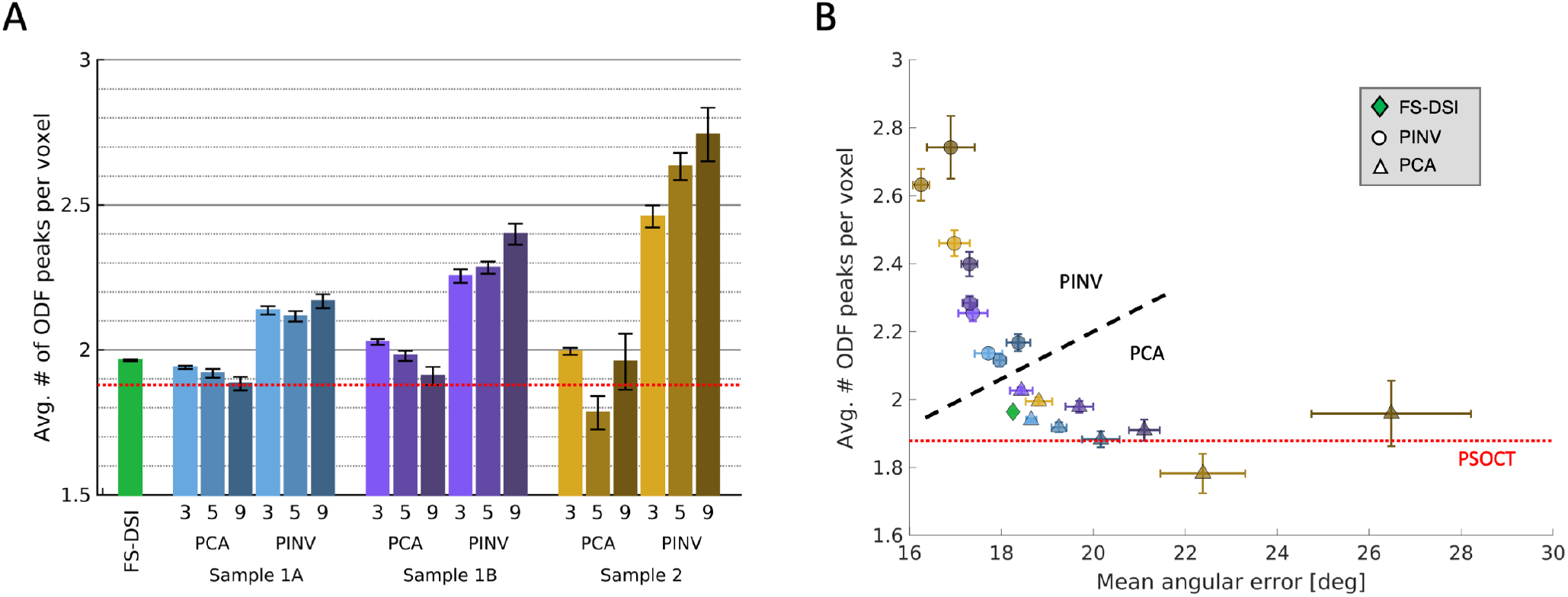
Number of reconstructed ODF peaks. (**A**) Average number of ODF peaks per WM voxel from fully sampled (green bar) and CS reconstructed (blue, purple, and brown bars) DSI data, and the average number of PSOCT FOD peaks per voxel (red dashed line). CS results show the average from the 9 volumes reconstructed with different CS undersampling masks. Error bars display the standard error. (**B**) Number of ODF peaks per voxel as a function of the mean angular error for fully sampled (green diamond) and CS reconstructed q-space data. The red dashed line indicates the average number of PSOCT FOD peaks per voxel. Circle markers correspond to PINV, and triangle markers correspond to PCA. Marker colors are the same as the bar plots in (A). Error bars indicate the standard error. PINV reconstructions (above black dashed line) produced more peaks and lower angular errors than PCA (below black dashed line). Most dMRI reconstructions produced more peaks per voxel than was observed in PSOCT. The CS-DSI reconstructions that were closest to FS-DSI on both axes were PCA with acceleration R=3.

FS-DSI produced an average of approximately 2 peaks per voxel, which was slightly greater than PSOCT (1.88 peaks per voxel). For an acceleration factor of R=3, PCA reconstructions produced a similar number of peaks compared to both PSOCT (less than 7.9% difference) and FS-DSI (less than 3.2% difference). As the acceleration factor increased, PCA tended to return slightly fewer peaks. On the other hand, all PINV reconstructions returned a notably greater number of peaks than both PSOCT (greater than 12.54% difference) and FS-DSI (greater than 7.6% difference). This number increased as the acceleration factor increased, and when the training data came from a different sample than the test data (*e.g.*, PINV with sample 2 training had greater than 30% difference vs. PSOCT for all acceleration factors).

Figure 5B plots the average number of peaks per voxel against the mean angular error. The PCA and PINV reconstructions are denoted by triangle and circle markers, respectively, and colored as in Figure 5A. Vertical and horizontal error bars indicate the standard error for each metric. FS-DSI (green diamond) served as the benchmark in terms of angular error. The red dotted line denotes the number of peaks per voxel for PSOCT. As indicated by the black dashed line (Figure 5B), there was a clear separation between PINV (above the line) and PCA (below the line). FS-DSI was situated at the knee of the curve, and PCA reconstructions with an acceleration factor of R=3 were closest to FS-DSI, regardless of the sample that was used as the training data set. PINV reconstructions displayed reduced angular errors, but also had significantly more peaks per voxel than PSOCT. Thus, we conclude that CS-DSI with a combination of R=3 acceleration and PCA reconstruction preserved the accuracy of FS-DSI with respect to the reference PSOCT orientations, but without introducing spurious peaks.

### 3.3. Effect of SNR on reconstruction error metrics

Figure 6A shows the slice-wise mean angular errors with respect to PSOCT, plotted against the SNR of the corresponding slice. Error bars indicate the standard error. The CS reconstructions included here used acceleration R=3 and dictionaries trained on high-SNR data from each sample. The curves for CS-DSI reconstructions closely resembled those of FS-DSI (green line) throughout the entire volume, with less than 2.52° variation in mean angular errors between all DSI reconstructions in each slice. FS-DTI (red line) had significantly greater errors than both FS-DSI and CS-DSI and showed a sharp increase in error as SNR decreased. The DSI angular errors were relatively robust to decreased SNR.

**Figure 6.**
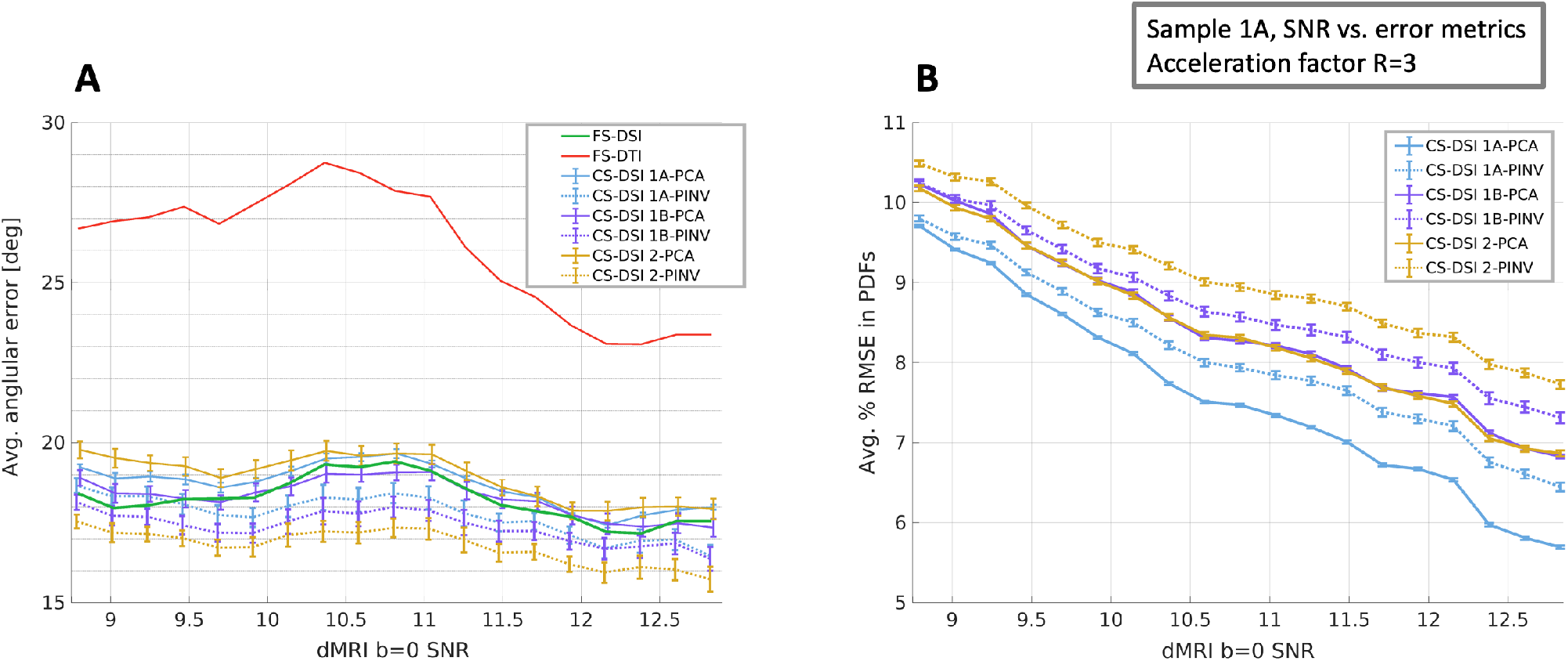
Reconstruction error as a function of SNR. **(A)** Average angular error of FS-DSI, FS-DTI, and CS-DSI with respect to reference axonal orientations from PSOCT. **(B)** Average RMSE in PDFs between CS-DSI and FS-DSI. Each error metric is averaged across all WM voxels in each slice and plotted against the SNR of the corresponding b=0 dMRI slice. Line colors and styles correspond to different dMRI reconstructions. For CS reconstructions, each error metric was averaged across 9 CS undersampling masks, with error bars showing the standard error of the mean across the 9 masks. For CS-DSI, line colors denote different training samples and line styles denote different CS reconstruction methods. All CS reconstructions used an acceleration factor of R=3 and high-SNR training data.

Table 3 shows statistics from the linear regressions of the mean angular error against SNR. The slopes were noticeably flatter for FS-DSI (−0.24° per unit SNR, p=0.051) and CS-DSI reconstructions (−0.28° to −0.46° per unit SNR, p<0.003) than FS-DTI (−1.20° per unit SNR, p=0.00011).

**Table 3.**
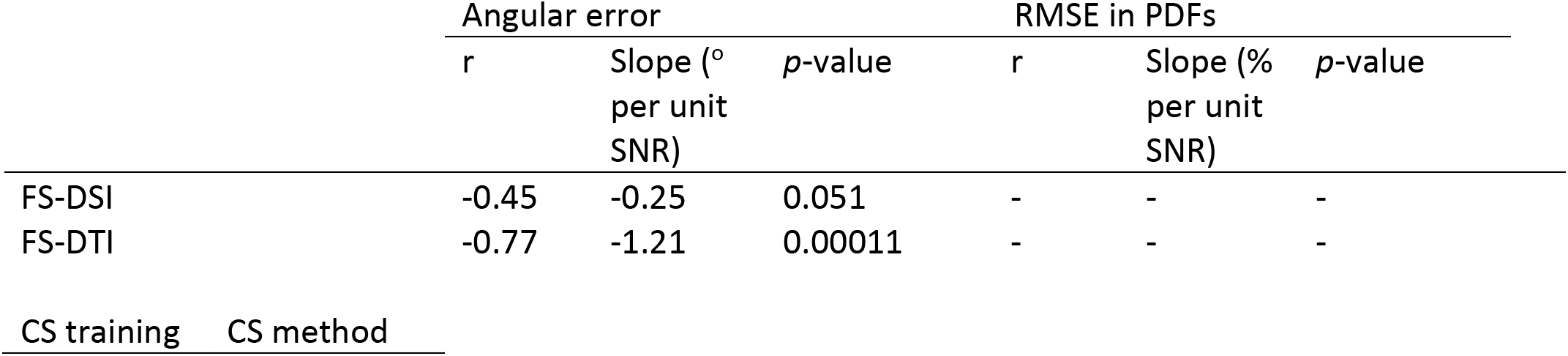
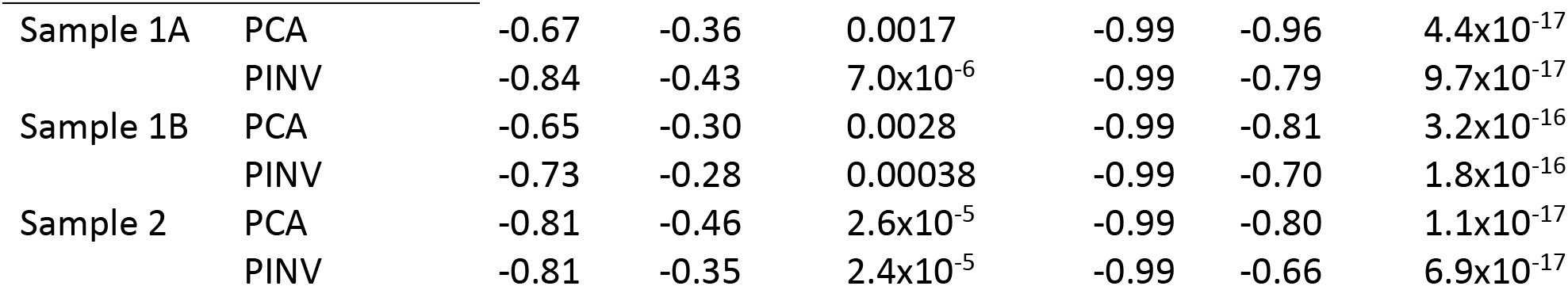
Linear regressions of reconstruction error metrics vs. SNR. Linear correlation coefficient (r), slope, and p-value from the linear regressions of the angular error with respect to PSOCT and the RMSE in PDFs with respect to fully sampled data as a function of SNR. CS reconstructions used an acceleration factor of R=3 and high-SNR training data.

|  | Angular error |  |  | RMSE in PDFs |  |  |
| --- | --- | --- | --- | --- | --- | --- |
| | $r$ | Slope ($^\circ$ per unit SNR) | $p$-value | $r$ | Slope (% per unit SNR) | $p$-value |
| FS-DSI | -0.45 | -0.25 | 0.051 | - | - | - |
| FS-DTI | -0.77 | -1.21 | 0.00011 | - | - | - |

| CS training | CS method |
| --- | --- |

**Table 3. Linear regressions of reconstruction error metrics vs.**
|  |  |  |  |  |  |  |  |
| --- | --- | --- | --- | --- | --- | --- | --- |
| Sample 1A | PCA | -0.67 | -0.36 | 0.0017 | -0.99 | -0.96 | $4.4 \times 10^{-17}$ |
| | PINV | -0.84 | -0.43 | $7.0 \times 10^{-6}$ | -0.99 | -0.79 | $9.7 \times 10^{-17}$ |
| Sample 1B | PCA | -0.65 | -0.30 | 0.0028 | -0.99 | -0.81 | $3.2 \times 10^{-16}$ |
| | PINV | -0.73 | -0.28 | 0.00038 | -0.99 | -0.70 | $1.8 \times 10^{-16}$ |
| Sample 2 | PCA | -0.81 | -0.46 | $2.6 \times 10^{-5}$ | -0.99 | -0.80 | $1.1 \times 10^{-17}$ |
| | PINV | -0.81 | -0.35 | $2.4 \times 10^{-5}$ | -0.99 | -0.66 | $6.9 \times 10^{-17}$ |

For comparison, Figure 6B shows the slice-wise mean RMSE in PDFs between each CS-DSI reconstruction and the fully sampled DSI data, as a function of SNR. The mean RMSE was calculated from the real part of the diffusion PDFs and averaged across all WM voxels in each slice. Error bars depict the standard error. Statistics from the linear regressions of the RMSE versus SNR are also given in Table 3. All reconstructions exhibited a strong negative correlation between RMSE and SNR (r=0.99, p<0.001) and showed a nearly linear increase in RMSE as SNR decreased (Figure 6B), with consistent linear regression slopes (−0.66 to −0.96 % RMSE per unit SNR). It should be noted that the linear fit of SNR likely smoothed the observed relationship, but that notwithstanding, the correlation between SNR and CS RMSE remained markedly apparent across all reconstructions.

### 3.4. Effect of SNR on dictionary training

After observing that CS reconstructions at acceleration R=3 using high-SNR training data performed nearly as well as FS-DSI in terms of angular error, we tested whether this was also true with low-SNR training data. The bar plot in Figure 7 compares the effect of training data SNR on the mean angular error of CS reconstructions at acceleration R=3 across all WM voxels analyzed. The CS reconstructions included here used one of the R=3 undersampling masks (out of the 9 undersampling masks used for Figures 4–6). All CS reconstructions exhibited greater mean angular error when using low-SNR than high-SNR training data, although the extent of differences varied depending on the training sample and reconstruction method. Specifically, PCA was more sensitive to the SNR level of the training data than PINV.

**Figure 7.**
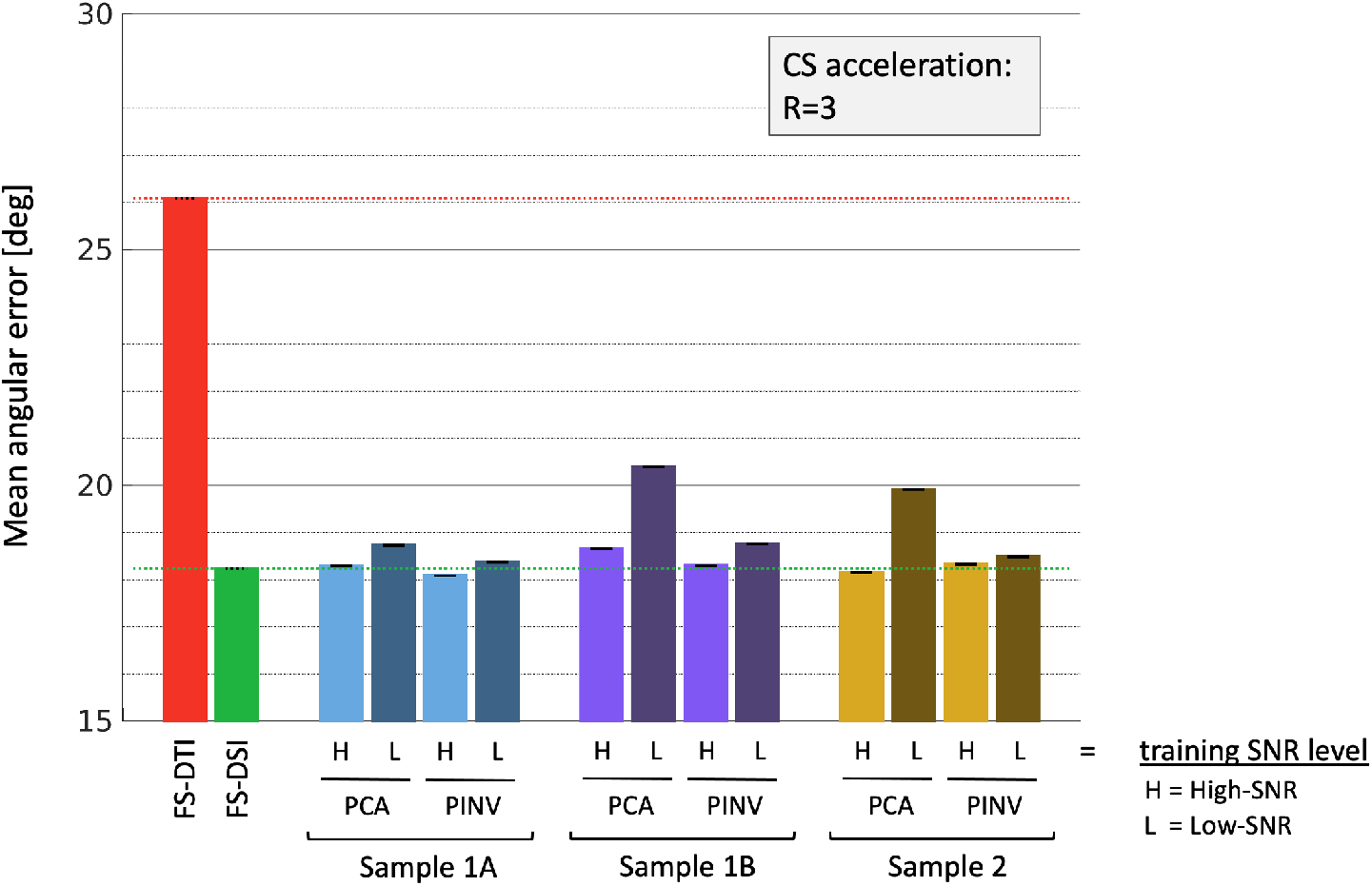
Effect of the SNR of the training data on angular error. Mean angular error across all analyzed WM voxels for CS reconstructions at acceleration R=3. Dictionaries were trained using either high-SNR (“H”, light shade) or low-SNR (“L”, dark shade) slices from sample 1A (blue), sample 1B (purple), or sample 2 (yellow). Results from FS-DTI (red) and FS-DSI (green) are shown on the far left. Error bars show standard error across voxels.

### 3.5. Accuracy of resampled q-shell data

Results from q-shell approximation are shown in Figure 8. The average relative NRMSE of each resampled diffusion-weighted volume with respect to the acquired shell data (Figure 8, top left) are plotted for NUFFT(FS) and NUFFT(CS) reconstructions (solid and dashed lines, respectively). The corresponding statistics are given in Table 4. NUFFT(FS) and NUFFT(CS) had nearly identical mean relative NRMSE for all shells (Table 4). The errors increased slightly as the b-value of the target shell increased, but were below 1 on average, for all shells. These results indicated that the resampling introduced negligible errors when compared to the variance between the two repeated acquisitions resulting from imaging noise. Additionally, resampling the undersampled q-space data without CS-DSI reconstruction yielded relative NRMSE values 2-3 fold larger than NUFFT(CS) (Supplemental Information; Figure S4, Table S1), indicating that dense grid q-space sampling, whether acquired with FS-DSI or recovered with CS-DSI, is essential for accurate q-shell approximation.

**Table 4.** Error of NUFFT q-space resampling. Relative NRMSE over white matter voxels from all directions of each q-shell.

| Shell b-value (s/mm<sup>2</sup>) | Relative NRMSE (mean +/- standard deviation) |  |
| --- | --- | --- |
|  | NUFFT(FS) | NUFFT(CS) |
| 4,000 | 0.78 +/- 0.073 | 0.78 +/- 0.064 |
| 12,000 | 0.83 +/- 0.11 | 0.83 +/- 0.11 |
| 20,000 | 0.97 +/- 0.27 | 0.95 +/- 0.29 |

**Figure 8.**
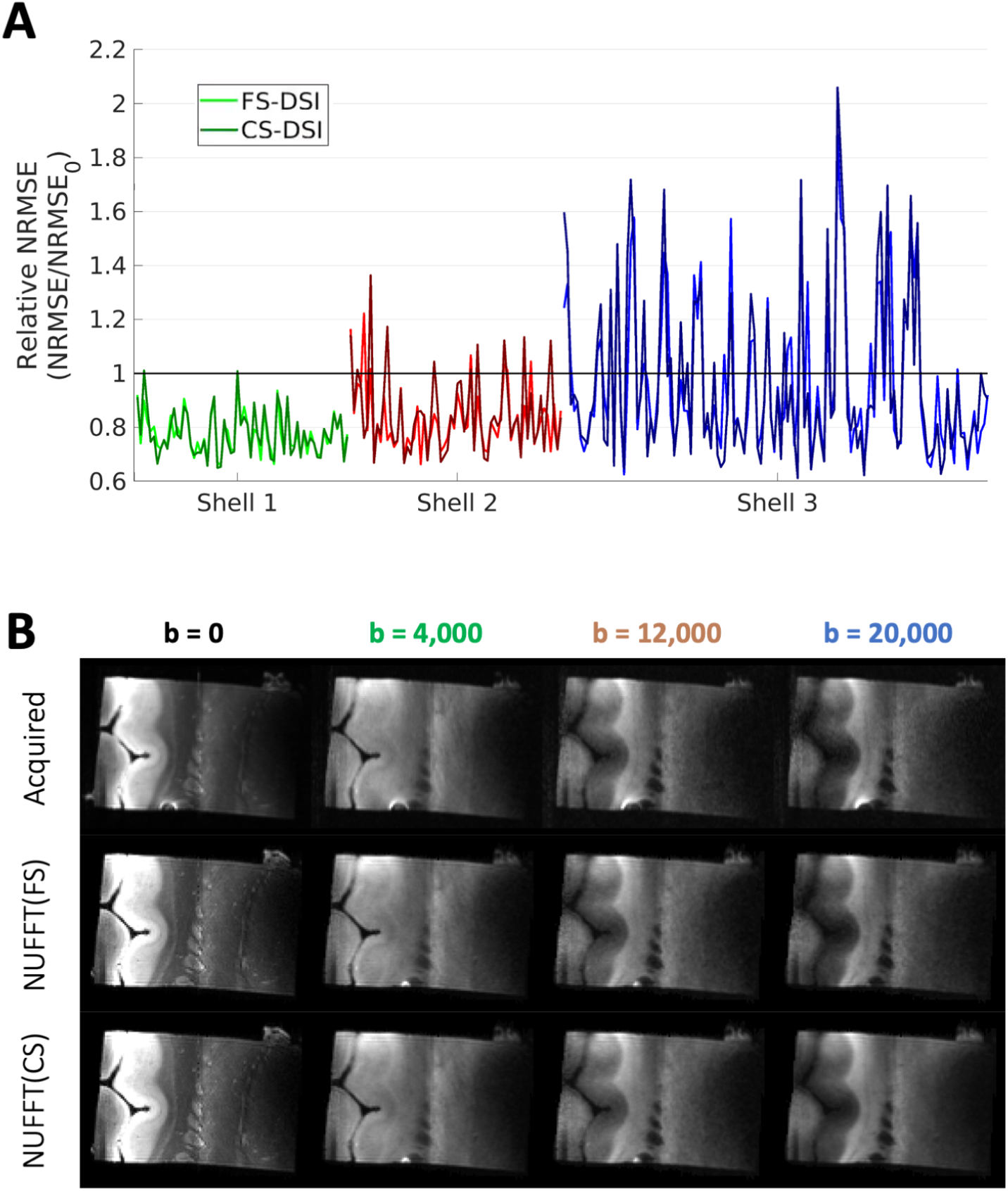
Results from NUFFT q-space resampling. (**A**) Relative normalized root-mean-square (NRMSE) from q-space resampling of FS-DSI (light shade) and CS-DSI (dark shade) grid q-space data for DWIs from the b=4,000 (green), 12,000 (red), and 20,000 s/mm^2^ (blue) shells. (**B**) Example slices of representative DWIs from acquired (top row), resampled FS-DSI (middle row) and resampled CS-DSI (bottom row) multi-shell data. The same grayscale display window was used for each column.

Figure 8B shows images of one slice from the b=0 volumes (left column) and one randomly selected DWI from each shell (three right-most columns). Both the NUFFT(FS) and NUFFT(CS) slices (middle and bottom rows, respectively) closely resembled the acquired multi-shell slices, illustrating that accurate resampling was possible using both FS-DSI and CS-DSI q-space data. Note that the b=0 images for the NUFFT(FS) and NUFFT(CS) data were the same.

Figure 9 displays microstructural maps obtained from DKI fitting of multi-shell data from sample 1B. The maps fitted using the NUFFT(FS) (middle row) and NUFFT(CS) (bottom row) data showed good agreement with the maps fitted using the acquired multi-shell data (top row). Kurtosis maps (MK, AK, RK; 3 right-most columns) were nearly identical between the acquired and NUFFT data. Minor increases in FA were observed in the lower half of the NUFFT(CS) map (second column from the left). All maps displayed similar features in regions of WM, as well as in subcortical (thalamus, right side of slice; putamen, bottom middle of slice) and cortical (left side of slice) GM regions.

**Figure 9.**
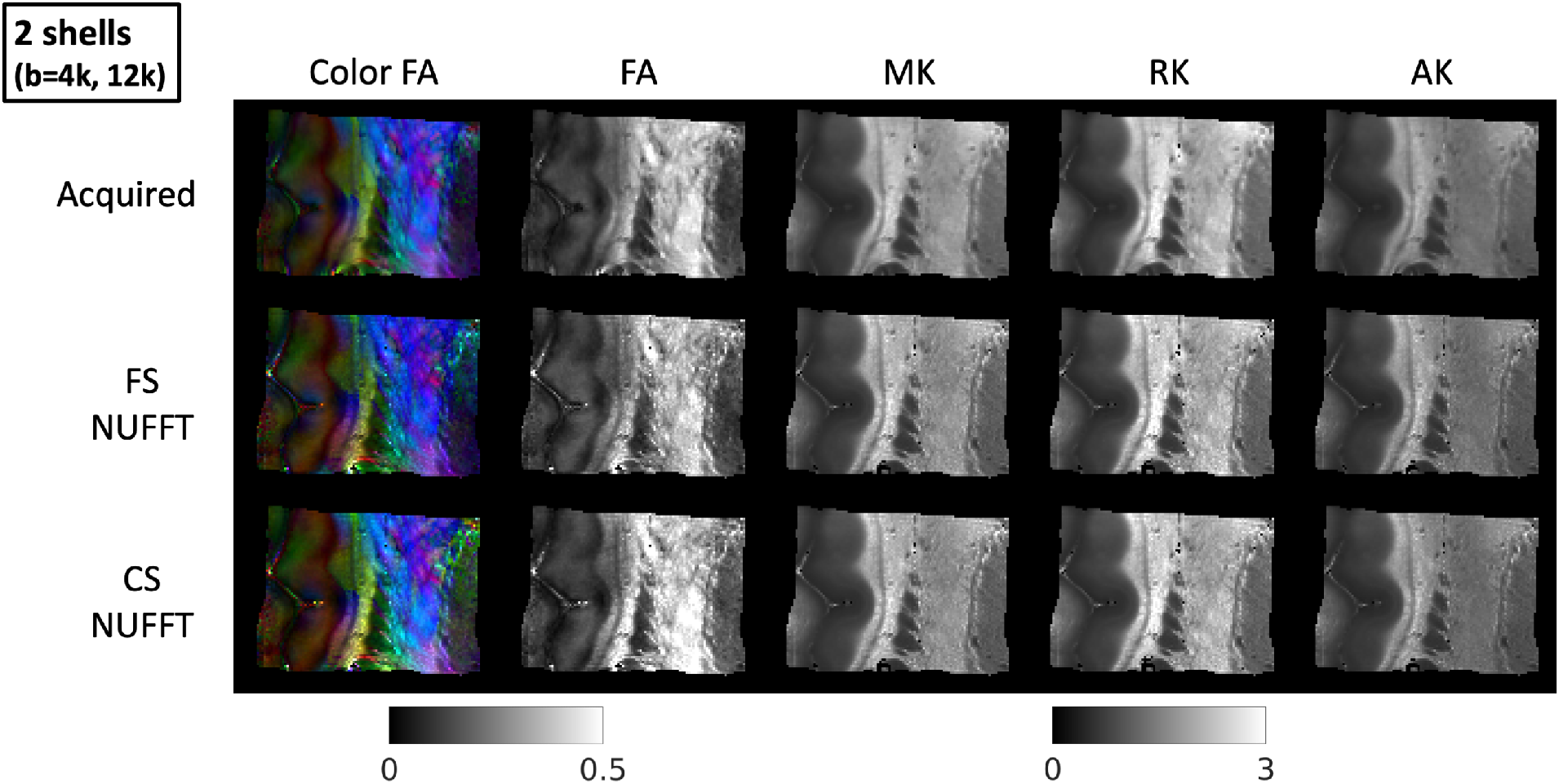
Microstructural maps extracted from multi-shell dMRI data. Representative slice showing color-encoded FA, FA, MK, RK and AK maps (from left to right, respectively) from DKI fitting of shells 1 and 2, for acquired multi-shell data (top row), FS-DSI data resampled onto shells (middle row), and CS-DSI data resampled onto shells (bottom row) data. Overall, the maps from resampled data exhibited good agreement with the maps fitted on the acquired data.

## 4. Discussion

In this study we performed post mortem validation of CS-DSI and its ability to approximate both grid- and shell-based diffusion sampling schemes. In one set of experiments, we evaluated the accuracy of diffusion orientation estimates obtained from CS-DSI by comparing them to microscopic-resolution reference measurements of fiber orientations from optical imaging. This is in contrast to previous *in* vivo validations that used simulated or fully sampled DSI data as the ground truth (Bilgic et al. 2013; Menzel et al. 2011; Paquette et al. 2015). Our results suggest that error metrics based on the difference between CS and fully sampled DSI in q-space may underestimate the accuracy of CS-DSI in capturing the underlying fiber geometry of the tissue. Specifically, although the difference in the PDFs obtained from CS and fully sampled DSI increases noticeably as the SNR decreases (Figure 6B), this translates to only a minor increase of the angular error of the peak orientations with respect to ground-truth measurements from optical imaging (Figure 6A). Note that these trends were observed both for reconstructions that tended to add spurious diffusion peaks (PINV) and for those that did not (PCA), hence the robustness of the angular error as a function of SNR did not appear to be explained by the number of peaks. The increasing error between PDFs obtained from CS and fully sampled DSI as SNR decreases is likely exacerbated by the fact that the fully sampled data are also corrupted by increasing levels of noise. This illustrated the importance of having objective measurements of the ground-truth anatomy from an independent modality, such as the optical imaging used here.

We employed two dictionary-based algorithms for CS reconstruction and investigated the effects of training data and undersampling rate on the accuracy of estimated fiber orientations. We found that PCA reconstruction at an acceleration factor of R=3 retained the fiber orientation accuracy of the fully sampled DSI data (Figure 4), without introducing spurious peaks (Figure 5). Its performance was as good when the training data for the dictionary came from a different sample than the test data as it was when the training and test data came from the same sample. However, we also found that it was important to use high-SNR training data to achieve this performance (Figure 7).

The PINV method achieved low angular errors at all acceleration factors and for all training samples (Figure 4). However, it did so at the cost of significant increases in the number of reconstructed peaks (Figure 5B), particularly at higher acceleration factors. Increases in the angular error and in the number of peaks can be thought of as losses of sensitivity and specificity, respectively. Therefore, both types of error are undesirable. Based on our findings, we recommend a CS acceleration factor of R=3 and PCA reconstruction, as this combination achieved both low angular error and low number of spurious peaks.

At an acceleration factor of R=3, the CS-DSI scheme comprises 171 diffusion-encoding gradient directions. This brings its acquisition time in line with those of state-of-the-art, multi-shell, high angular resolution diffusion imaging (HARDI) acquisitions. Our findings suggest that CS-DSI allows us to approximate both a fully sampled, 514-direction DSI scheme (via CS reconstruction) and multi-shell schemes (via q-space resampling). Traditionally, HARDI vs. DSI has been framed as a choice between two mutually exclusive approaches, but this does not have to be the case. We have previously shown that fully sampled DSI data can be resampled onto q-shells (Jones et al. 2020). Here we show that an accelerated CS-DSI scheme also allows resampling onto arbitrary q-shells (of course, to within the limits placed by the bmax of the DSI acquisition). We find that both the q-space approximation error (Table 4, Figure 8) and the microstructural measures obtained after resampling the data onto q-shells (Figure 9) are comparable between CS and fully sampled DSI data.

The findings of recent validation studies suggest that DSI may provide more accurate fiber reconstructions than single- or multi-shell acquisition schemes. DSI produced more accurate fiber orientation estimates in simulations (Daducci et al. 2013) and comparisons to optical imaging measurements (Jones et al. 2020), as well as more accurate tractography when compared to ground-truth anatomic tracing in non-human primates (Maffei et al. 2021, 2020). The present study shows that a sparsely sampled CS-DSI protocol preserves the high angular accuracy of fully sampled DSI. It also preserves the flexibility of resampling the data onto q-shells at arbitrary b-values, to facilitate analyses that require shelled data, such as DKI (Jensen et al. 2005), neurite orientation dispersion and density imaging (Zhang et al. 2012), etc. At the same time, it allows direct reconstruction of the full EAP. This allows a wider range of analyses to be performed. For example, diffusion PDFs have been previously used to characterize age-related WM demyelination (Fatima et al. 2013), delineate pathological tissue lesions in patients with multiple sclerosis (Assaf et al. 2002), and map *in vivo* axon caliber in the human brain (Hori et al. 2016). Furthermore, we have recently shown that diffusion EAPs may be used to improve the decision making of tractography algorithms (A. Yendiki et al. 2020) or reconstruct generalized anisotropy profiles that provide contrast between different gray- and white-matter structures (Jones et al. 2021).

In regard to the dictionary-based CS algorithms that we investigated, our findings indicate that dictionary generalizability differs between the PINV and PCA methods, and that the extent of these differences depends on both the CS acceleration factor (Figure 4) and the training data SNR (Figure 7). One potential factor contributing to these differences may be the ways in which PDFs are represented within the algorithms. PCA performs reconstructions in a reduced-dimensionality space consisting of only the *T* principle components that describe the greatest variance in the training PDFs, and the optimal number of components *T* decreases at higher acceleration factors in order to improve the conditioning of the pseudoinverse in the least-squares reconstruction (Bilgic et al. 2013). In our experiments, *T* was around 20 at acceleration R=3 and less than that at higher accelerations, which are moderately lower than the optimal *in vivo* parameters. Intuitively, reducing the number of principal components subsequently limits the ability to describe finer scale details in PDFs, and, together with the extremely undersampled q-space data used at high accelerations, likely hinders the level of detail in such reconstructions. In contrast, PINV operates directly on the training PDFs themselves, exploiting the prior information encoded in the dictionary atoms to bypass sparsity constraints. Together, these differences may contribute to the different behavior of PINV and PCA as the acceleration factor increases.

The PINV method investigated here uses a dictionary containing PDFs from a slice of fully sampled training data, without any further training. An alternative approach is to use a dictionary trained with the K-SVD algorithm (Aharon, Elad, and Bruckstein 2006), which enhances the sparsity level of PDF representations and is a fraction of the size of the PINV dictionary, allowing up to a 50% reduction in computation time. Regardless of the dictionary, reconstructions are performed using the Tikhonov-regularized pseudoinverse. Previous comparisons between PINV using a 3191-column dictionary and PINV(K-SVD) using a 258-column dictionary reported nearly identical reconstruction quality in terms of RMSE, as well as equivalent representational power between the two dictionaries (Bilgic et al. 2013). Although we did not include results from PINV(K-SVD) here, we did perform CS reconstructions and angular error analysis for PINV(K-SVD). We observed very similar results to PINV, both in terms of RMSE with respect to fully sampled data and in terms of angular error with respect to PSOCT. It is possible that the behavior of the PINV reconstruction that we observed in these experiments was related to the optimization of the Tikhonov regularization parameter λ. The optimal λ was selected to minimize the RMSE in PDFs with respect to FS-DSI, which may have been sub-optimal in terms of spurious peaks.

### Relation to previous studies

We have previously used the present PSOCT analysis framework (and sample 1A from this work) in an extensive validation study (Jones et al. 2020). In that work, we assessed the accuracy of fiber orientations estimated from various dMRI orientation reconstruction methods and sampling schemes, including DSI, single-, and multi-shell. The DSI results are somewhat different between the two studies because the DSI ODF reconstruction is different. In (Jones et al. 2020), DSI reconstructions were performed with the DSI Studio toolbox (http://dsi-studio.labsolver.org), and used filtered q-space signals (Hanning window, width=16) with default parameters (*e.g.*, zero-padded 16×16×16 q-space grid, ODF integration lower/upper bounds of 0.25x and 0.75x FOV), and a ODF peak threshold of 0.01 (1%). Here, we used a different approach for ODF reconstruction, imposed a (slightly) more stringent peak threshold (5%) and included a peak separation threshold. These factors likely contributed to the ~1-2° increases in mean angular error reported here compared to previous results. Nonetheless, the angular errors in this work were similar to the best performing reconstructions in (Jones et al. 2020), namely DSI and generalized q-sampling imaging (GQI) (Yeh, Wedeen, and Tseng 2010) with the fully sampled DSI data and q-ball imaging (QBI) (Tuch 2004; Aganj et al. 2010) with single- and multi-shell data, which were between ~17° and ~19°.

This work analyzed the dictionary-based CS-DSI methods introduced by Bilgic et. al. (2013), however there were several technical differences between these works that should be noted. First, in terms of data acquisition, we used DSI data from *ex vivo* human brain samples acquired at 9.4T with a 4-channel surface Rx coil, whereas Bilgic et. al. used *in vivo* DSI from the 3T Connectom system with a custom 64-channel head coil (Keil et al. 2013). These differences did not have an apparent effect on the dictionary-based CS-DSI methods. First, the CS algorithms yielded similar results when using *ex vivo* training and test data as when using *in vivo* training and test data. Whether the same would also be true when using *ex vivo* training data and *in vivo* test data (or vice versa) has yet to be investigated. Such an approach may be of interest as long, *ex vivo* acquisitions can be a way to collect very high-SNR training data. Second, while both studies generated dictionaries with PDFs from a single slice of fully sampled data, a single slice of our *ex vivo* samples covers only a small anatomical region, whereas a slice of *in vivo* data covers an entire cross-section of the brain. Given that our training samples were cut from different anatomical locations, one might expect that local microstructural differences between training and test samples might pose challenges for dictionary generalizability. However, our findings showed that high-quality reconstructions could be obtained using training and test data from different samples (Figure 7), indicating that our *ex vivo* dictionaries possess the representational power to generalize across samples. Indeed, the “residual” (Bilgic et al. 2013) between *ex vivo* PDF dictionaries, *i.e.*, the energy of the part of one dictionary that cannot be represented by another, was negligibly small (~10^−12^), confirming that dictionaries from different samples possess equivalent representational power.

### Limitations

The present study did not evaluate all possible methods for CS-DSI reconstruction. However, it included one method that achieved good performance on all metrics (PCA on R=3), and highlighted the two different ways in which performance can degrade as the acceleration factor increases (Figure 5B). The PCA method exhibited diminishing performance in terms of angular error, a measure of sensitivity, while the PINV method exhibited significant increases in peaks per voxel, a decrease in specificity (Figure 5B). These results could be used as a benchmark for future CS-DSI methods, as well as for similar deep learning-based approaches for reconstructing sparse q-space samplings (Golkov et al. 2016; Gibbons et al. 2019). These approaches may allow even more efficient acceleration and preserve accuracy at acceleration factors greater than R=3.

The two dictionary-based CS-DSI reconstruction methods that we evaluated here use discrete EAP representations and L2-regularized algorithms. We studied these methods because they are fast, reconstructing an entire slice in a matter of seconds, and easy to implement, using dictionaries of fully sampled PDFs without any additional training, and having only one free parameter to determine. However, there are various other CS-DSI methodologies that utilize other basis functions (*e.g.*, discrete cosine transform, discrete wavelet transform) or approaches to solving the underdetermined CS problem (*e.g.*, L1-regularized methods such as equality constrained or regularized Dictionary-FOCUSS). While the L2-regularized dictionary-based CS-DSI methods investigated here were shown to provide comparable reconstruction quality to both fixed transforms and iterative L1-regularized methods (Bilgic et al. 2013), those evaluations were mostly based on RMSE with respect to fully sampled DSI data. Analyzing the fidelity of other CS-DSI algorithms in the framework presented here could be valuable, but the lengthy computation times accompanying iterative algorithms would ultimately limit their utility. We opted to focus our investigation on the two dictionary-based CS methods, PCA and PINV, which were computationally tractable and thus enabled us to thoroughly study the effects of experimental factors on their performance.

In this work, we used PSOCT to obtain ground-truth measurements of fiber orientations. There are several other techniques that have been used to assess the accuracy of dMRI orientation estimates. For a comprehensive discussion of their relative merits and weaknesses, we refer the reader to a recent review (Anastasia Yendiki et al. 2021). One approach is to extract orientations from myelin-stained sections (Leergaard et al. 2010; Choe et al. 2012; K. Schilling et al. 2017; Seehaus et al. 2015) or from confocal microscopy of slices stained with DiI, a fluorescent dye (Budde and Frank 2012). Quantification of 3D orientations has been reported with Dil stained slices (K.G. Schilling et al. 2018; K. Schilling et al. 2016; Khan et al. 2015). When using histological stains, the fiber orientation angles have to be obtained either by manual tracing or by an image processing step such as structure-tensor analysis. This step may introduce a source of error.

Optical imaging based on light polarization is a *label-free* approach that provides direct measurements of fiber orientations by exploiting the intrinsic optical property of tissue birefringence. In addition to PSOCT, polarized light imaging (PLI) also uses birefringence to measure axonal orientations (Mollink et al. 2017; Henssen et al. 2019; Axer et al. 2011). However, unlike PSOCT, PLI requires tissue to be sectioned and mounted before imaging. This can lead to severe tissue distortions that demand a complex registration framework to correct (Majka and Wójcik 2016; Ali et al. 2017; Ali et al. 2018). PSOCT images the blockface of tissue before slicing, greatly reducing tissue distortions and allowing accurate volumetric reconstructions.

The PSOCT technique has its own limitations. Notably, the optic axis orientation measurements do not describe the 3D orientation, but rather its projection onto the imaging plane. Fibers oriented orthogonal to the imaging plane have small in-plane components and may introduce uncertainty into the PSOCT optic axis measurements. Such fibers would also exhibit low retardance. To avoid biases from through-plane fibers, we only analyzed WM voxels with high retardance. In a previous study using an identical procedure, we showed that the in-plane angular errors between dMRI and PSOCT were consistently between 10° and 20°, regardless of the through-plane component of the dMRI orientations (Jones et al. 2020). This suggests that our in-plane angular errors are not biased by the presence of through-plane diffusion. Finally, the 2D angular errors in our PSOCT studies agree with both the 2D and 3D angular errors reported in studies that used histological staining, further supporting the validity of in-plane angular errors as a measure of accuracy (Anastasia Yendiki et al. 2021).

There are ways to bypass the 2D nature of the PSOCT optic axis measurements to interrogate 3D orientations. One possibility is to apply structure tensor analysis to PSOCT volumetric intensity data (Wang, Lenglet, and Akkin 2015; Wang et al. 2011). Alternatively, 3D fiber orientations can be obtained by collecting PSOCT optic axis measurements with multiple light incidence angles on the tissue surface and using these measurements to infer the through-plane orientation. This approach has been previously demonstrated in biological tissue (Nadya Ugryumova et al. 2009; Nadezhda Ugryumova, Gangnus, and Matcher 2006; Liu et al. 2016).

## 5. Conclusion

We have demonstrated that, when utilized in an appropriate manner, dictionary-based CS-DSI reconstructions can reduce acquisition times by a factor of 3 while preserving the accuracy of DSI fiber orientation estimates with respect to PSOCT. In particular, given an adequate SNR level of the training data, the PCA method produced high-fidelity reconstructions that reliably maintained the angular accuracy of fully sampled DSI data, without introducing spurious peaks. We also demonstrated that we could tolerate a sizeable increase in the RMSE between PDFs obtained from CS-DSI and fully sampled DSI data, without incurring a large decrease in the accuracy of the peak orientations with respect to the axonal orientations measured with optical imaging. This underscores the importance of having access to ground-truth measurements of fiber architecture from a modality that is independent of water diffusion and MRI measurement noise. Finally, we showed that a sparsely sampled CS-DSI acquisition, combined with q-space resampling, could be used to approximate not only fully sampled DSI but also multi-shell data with high accuracy, while keeping the acquisition time short. Our findings confirm the viability of CS-DSI as a technique for accelerating DSI acquisitions, while permitting a wide range of analyses that require either grid- or shell-based dMRI data. They also provide useful benchmarks for future development of undersampled q-space acquisition and reconstruction techniques.

## Acknowledgements

This work was funded by NIH grants R01-EB021265, K99-EB023993, R01-AG057672, R01-EB019956, R01-EB017337, U01-EB025162, P41-EB030006, R03-EB031175, and U01-MH117023. It was carried out at the Athinoula A. Martinos Center for Biomedical Imaging at the Massachusetts General Hospital, using resources provided by the Center for Functional Neuroimaging Technologies, P41-EB015896, a P41 Biotechnology Resource Grant supported by the National Institute of Biomedical Imaging and Bioengineering (NIBIB), National Institutes of Health. This work also involved the use of instrumentation supported by the NIH Shared Instrumentation Grant Program (S10RR025563, S10RR023401, S10RR019307, and S10RR023043).

## Abbreviations

dMRI: diffusion magnetic resonance imaging
DWI: diffusion weighted image
DSI: diffusion spectrum imaging
DTI: diffusion tensor imaging
PDF: probability density function
EAP: ensemble average propagator
PSOCT: polarization sensitive optical coherence tomography
SNR: signal-to-noise ratio
CS: compressed sensing
FS: fully sampled
PGSE: pulsed gradient spin echo
EPI: echo planar imaging
FT: Fourier transform
HARDI: high angular resolution diffusion imaging
ODF: orientation distribution function
WM: white matter
GM: gray matter
ROI: region of interest
GQI: generalized q-sampling imaging
QBI: q-ball imaging
PCA: principal component analysis
PINV: pseudoinverse
NUFFT: nonuniform fast Fourier transform
RMSE: root mean square error
FA: fractional anisotropy
FOCUSS: FOCal Underdetermined System Solver
SD: spherical deconvolution
G_max_: maximum gradient amplitude

## Supplemental data

**Figure S1.**
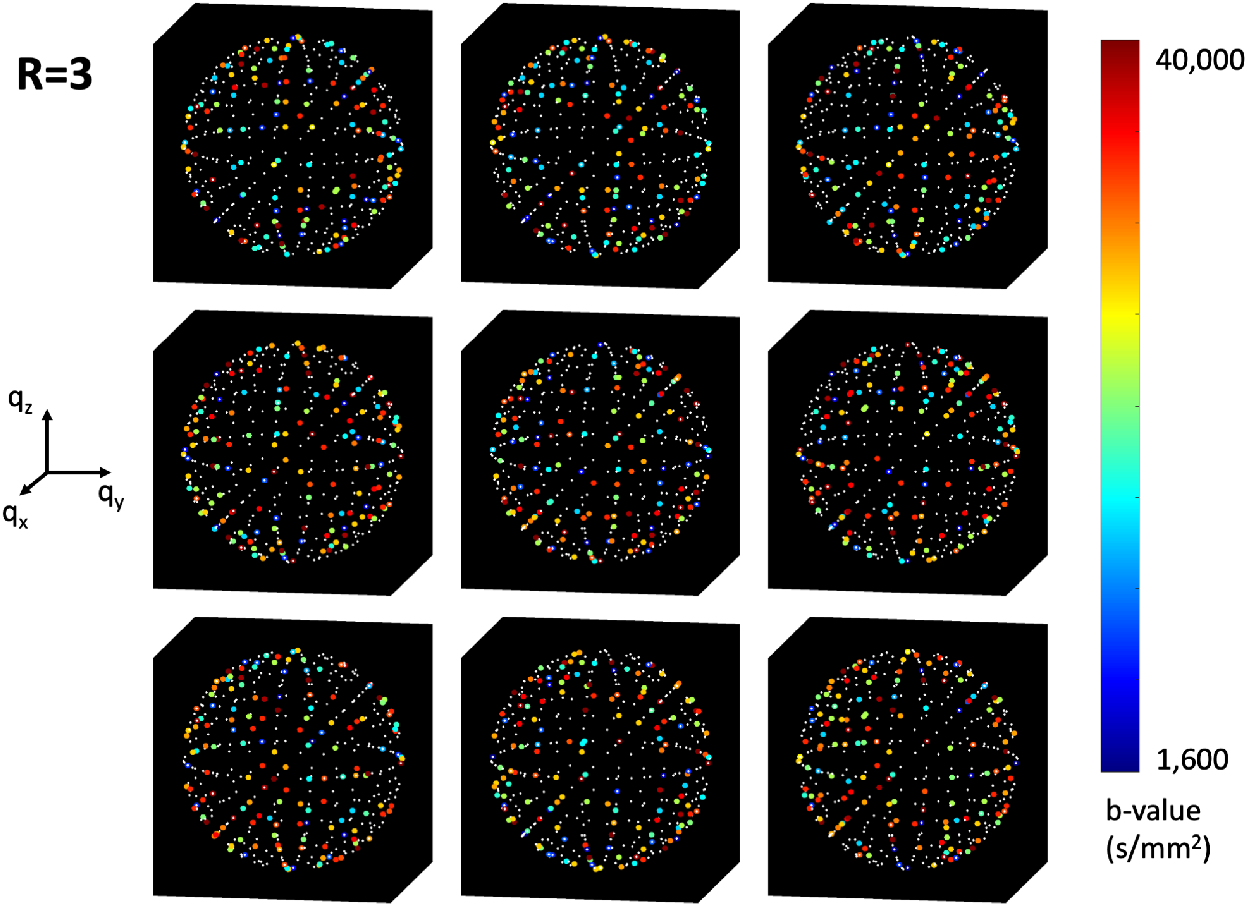
CS-DSI angular sampling patterns for acceleration R=3. Scatter plots of the nine different q-space undersampling masks, projected onto the unit sphere. Samples included in the mask are displayed as colored dots, with the color corresponding to b-value (see colorbar on far right). Samples not included in the mask (*i.e.*, that were reconstructed) are plotted as white dots.

**Figure S2.**
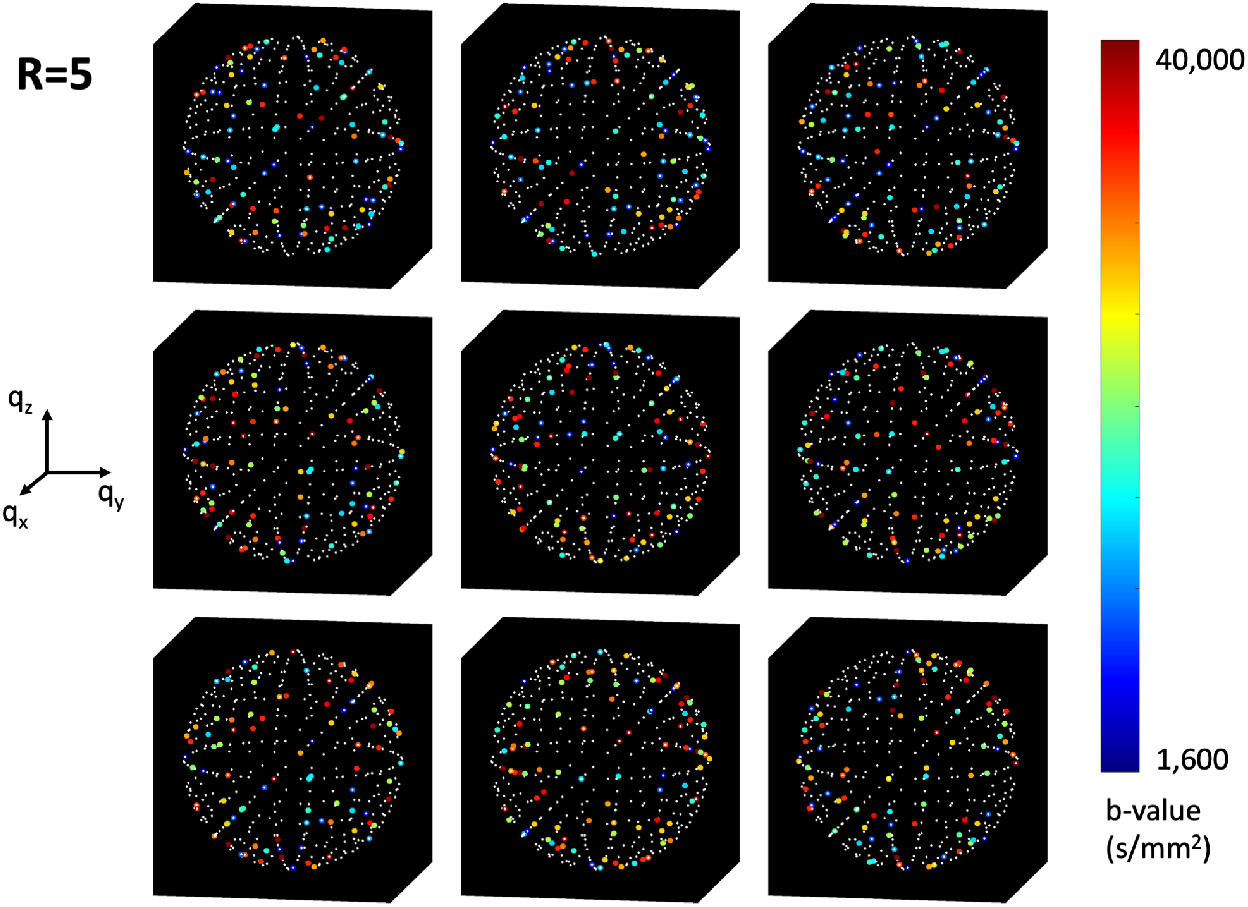
CS-DSI angular sampling patterns for acceleration R=5. Scatter plots of the nine different q-space undersampling masks, projected onto the unit sphere. Samples included in the mask are displayed as colored dots, with the color corresponding to b-value (see colorbar on far right). Samples not included in the mask (*i.e.*, that were reconstructed) are plotted as white dots.

**Figure S3.**
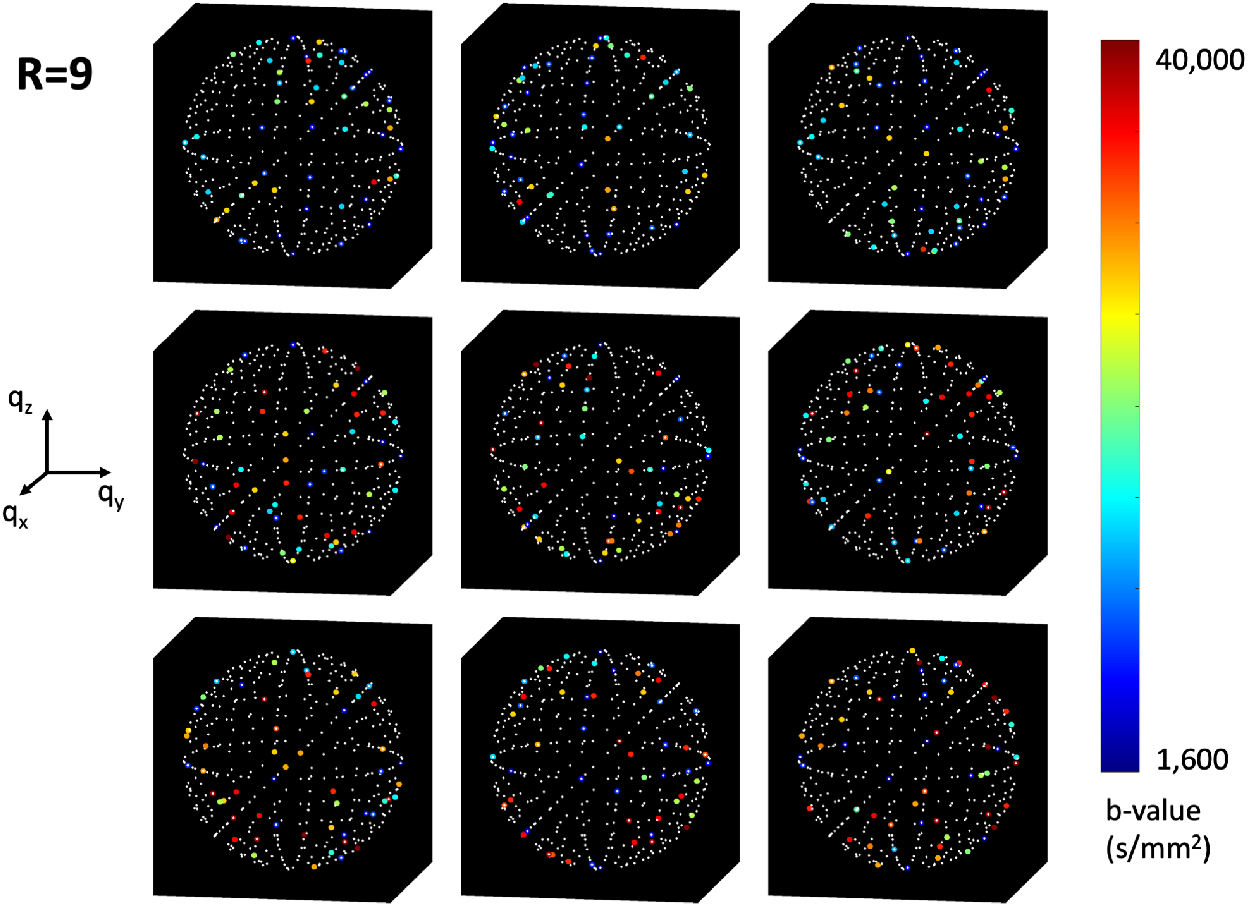
CS-DSI angular sampling patterns for acceleration R=9. Scatter plots of the nine different q-space undersampling masks, projected onto the unit sphere. Samples included in the mask are displayed as colored dots, with the color corresponding to b-value (see colorbar on far right). Samples not included in the mask (*i.e.*, that were reconstructed) are plotted as white dots.

**Figure S4.**
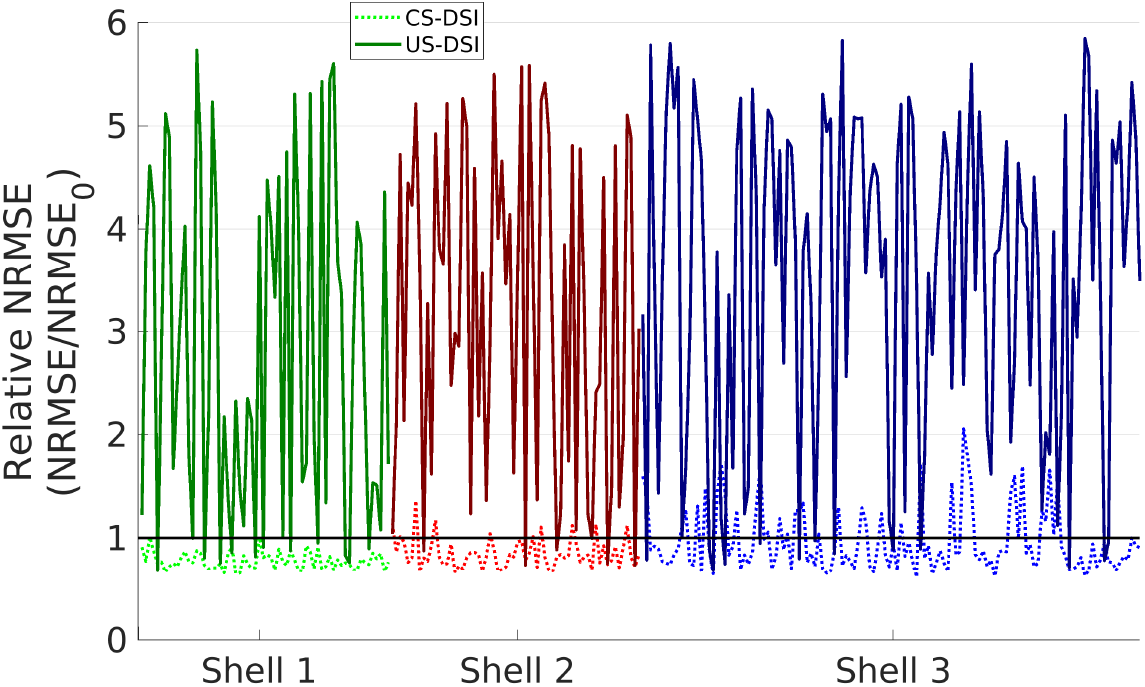
Results from NUFFT q-space resampling of undersampled vs CS-DSI q-space. Relative normalized root-mean-square (NRMSE) from q-space resampling of undersampled DSI q-space (US-DSI; solid line, dark shade) and CS-DSI (dashed line, light shade) grid q-space data at acceleration R=3 for DWIs from the b=4,000 (green), 12,000 (red), and 20,000 s/mm^2^ (blue) shells. The US-DSI data consisted of only the 171 q-space samples included in the undersampling mask, whereas the CS-DSI data also contained the additional grid samples reconstructed by CS.

**Table S1.**
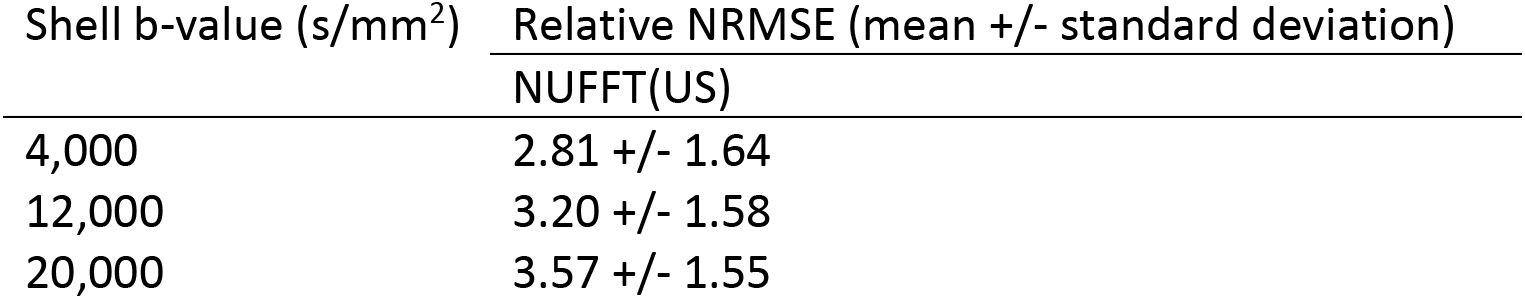
Error of q-space resampling from undersampled CS data. Relative NRMSE of data that were resampled on q-shells by applying the NUFFT directly to R=3 (171-direction) undersampled data. Mean and standard deviation are over WM voxels from all directions of each q-shell.

## Notes

### Competing Interest Statement

The authors have declared no competing interest.

## References

Aganj, Iman, Christophe Lenglet, Guillermo Sapiro, Essa Yacoub, Kamil Ugurbil, and Noam Harel. 2010. “Reconstruction of the orientation distribution function in single–and multiple– shell q–ball imaging within constant solid angle.” Magnetic resonance in medicine 64 (2): 554–566.

Aharon, Michal, Michael Elad, and Alfred Bruckstein. 2006. “K-SVD: An algorithm for designing overcomplete dictionaries for sparse representation.” IEEE Transactions on signal processing 54 (11): 4311–4322.

Ali, Sharib, Karl Rohr, Markus Axer, Katrin Amunts, Roland Eils, and Stefan Wörz. 2017. “Registration of ultra-high resolution 3D PLI data of human brain sections to their corresponding high-resolution counterpart.” 2017 IEEE 14th International Symposium on Biomedical Imaging (ISBI 2017).

Ali, Sharib, Stefan Wörz, Katrin Amunts, Roland Eils, Markus Axer, and Karl Rohr. 2018. “Rigid and non-rigid registration of polarized light imaging data for 3D reconstruction of the temporal lobe of the human brain at micrometer resolution.” NeuroImage 181: 235–251.

Assaf, Yaniv, D Ben-Bashat, J Chapman, S Peled, IE Biton, M Kafri, Y Segev, T Hendler, AD Korczyn, and M Graif. 2002. “High b–value q–space analyzed diffusion–weighted MRI: application to multiple sclerosis.” Magnetic Resonance in Medicine: An Official Journal of the International Society for Magnetic Resonance in Medicine 47 (1): 115–126.

Axer, Hubertus, Sindy Beck, Markus Axer, Friederike Schuchardt, Jörn Heepe, Anja Flücken, Martina Axer, Andreas Prescher, and Otto W Witte. 2011. “Microstructural analysis of human white matter architecture using polarized light imaging: views from neuroanatomy.” Frontiers in neuroinformatics 5: 28.

Basser, Peter J, James Mattiello, and Denis LeBihan. 1994a. “Estimation of the effective self-diffusion tensor from the NMR spin echo.” Journal of Magnetic Resonance, Series B 103 (3): 247–254.

Basser, Peter J, James Mattiello, and Denis LeBihan. 1994b. “MR diffusion tensor spectroscopy and imaging.” Biophysical journal 66 (1): 259–267.

Bilgic, Berkin, Itthi Chatnuntawech, Kawin Setsompop, Stephen F Cauley, Anastasia Yendiki, Lawrence L Wald, and Elfar Adalsteinsson. 2013. “Fast dictionary-based reconstruction for diffusion spectrum imaging.” IEEE transactions on medical imaging 32 (11): 2022–2033.

Bilgic, Berkin, Kawin Setsompop, Julien Cohen-Adad, Anastasia Yendiki, Lawrence L Wald, and Elfar Adalsteinsson. 2012. “Accelerated diffusion spectrum imaging with compressed sensing using adaptive dictionaries.” Magnetic Resonance in Medicine 68 (6): 1747–1754.

Boas, David A, Hui Wang, Caroline Magnain, and Bruce Fischl. 2017. “Polarization-sensitive optical coherence tomography of the human brain connectome.” SPIE Newsroom 10 (2.1201701): 006834.

Budde, Matthew D, and Joseph A Frank. 2012. “Examining brain microstructure using structure tensor analysis of histological sections.” Neuroimage 63 (1): 1–10.

Choe, AS, I Stepniewska, DC Colvin, Z Ding, and AW Anderson. 2012. “Validation of diffusion tensor MRI in the central nervous system using light microscopy: quantitative comparison of fiber properties.” NMR in Biomedicine 25 (7): 900–908.

Daducci, Alessandro, Erick Jorge Canales-Rodrı, Maxime Descoteaux, Eleftherios Garyfallidis, Yaniv Gur, Ying-Chia Lin, Merry Mani, Sylvain Merlet, Michael Paquette, and Alonso Ramirez-Manzanares. 2013. “Quantitative comparison of reconstruction methods for intra-voxel fiber recovery from diffusion MRI.” IEEE transactions on medical imaging 33 (2): 384–399.

De Boer, Johannes F, Christoph K Hitzenberger, and Yoshiaki Yasuno. 2017. “Polarization sensitive optical coherence tomography–a review.” Biomedical optics express 8 (3): 1838–1873.

De Boer, Johannes F, Thomas E Milner, Martin JC van Gemert, and J Stuart Nelson. 1997. “Two-dimensional birefringence imaging in biological tissue by polarization-sensitive optical coherence tomography.” Optics letters 22 (12): 934–936.

Donoho, David L. 2006. “Compressed sensing.” IEEE Transactions on information theory 52 (4): 1289–1306.

Fan, Chuanmao, and Gang Yao. 2013. “Imaging myocardial fiber orientation using polarization sensitive optical coherence tomography.” Biomedical optics express 4 (3): 460–465.

Fatima, Zareen, Utaroh Motosugi, Masaaki Hori, Toshiyuki Onodera, Keiichi Ishigame, Kazuo Yagi, and Tsutomu Araki. 2013. “Age-related white matter changes in high b-value q-space diffusion-weighted imaging.” Neuroradiology 55 (3): 253–259. https://doi.org/10.1007/s00234-012-1099-4.

Fessler, Jeffrey A, and Bradley P Sutton. 2003. “Nonuniform fast Fourier transforms using min-max interpolation.” IEEE transactions on signal processing 51 (2): 560–574.

Garyfallidis, Eleftherios, Matthew Brett, Bagrat Amirbekian, Ariel Rokem, Stefan Van Der Walt, Maxime Descoteaux, and Ian Nimmo-Smith. 2014. “Dipy, a library for the analysis of diffusion MRI data.” Frontiers in neuroinformatics 8: 8.

Gibbons, Eric K, Kyler K Hodgson, Akshay S Chaudhari, Lorie G Richards, Jennifer J Majersik, Ganesh Adluru, and Edward VR DiBella. 2019. “Simultaneous NODDI and GFA parameter map generation from subsampled q–space imaging using deep learning.” Magnetic resonance in medicine 81 (4): 2399–2411.

Golkov, Vladimir, Alexey Dosovitskiy, Jonathan I Sperl, Marion I Menzel, Michael Czisch, Philipp Sämann, Thomas Brox, and Daniel Cremers. 2016. “Q-space deep learning: twelve-fold shorter and model-free diffusion MRI scans.” IEEE transactions on medical imaging 35 (5): 1344–1351.

Gorodnitsky, Irina F, and Bhaskar D Rao. 1997. “Sparse signal reconstruction from limited data using FOCUSS: A re-weighted minimum norm algorithm.” IEEE Transactions on signal processing 45 (3): 600–616.

Guo, Shuguang, Jun Zhang, Lei Wang, J Stuart Nelson, and Zhongping Chen. 2004. “Depth-resolved birefringence and differential optical axis orientation measurements with fiber-based polarization-sensitive optical coherence tomography.” Optics Letters 29 (17): 2025–2027.

Hagmann, Patric, Leila Cammoun, Xavier Gigandet, Reto Meuli, Christopher J Honey, Van J Wedeen, and Olaf Sporns. 2008. “Mapping the structural core of human cerebral cortex.” PLoS biology 6 (7).

Henssen, Dylan JHA, Jeroen Mollink, Erkan Kurt, Robert van Dongen, Ronald HMA Bartels, David Gräβel, Tamas Kozicz, Markus Axer, and Anne-Marie Van Cappellen van Walsum. 2019. “Ex vivo visualization of the trigeminal pathways in the human brainstem using 11.7 T diffusion MRI combined with microscopy polarized light imaging.” Brain Structure and Function 224 (1): 159–170.

Hori, Masaaki, Kouhei Kamiya, Atsushi Nakanishi, Issei Fukunaga, Masakazu Miyajima, Madoka Nakajima, Michimasa Suzuki, Yuriko Suzuki, Ryusuke Irie, Koji Kamagata, Hajime Arai, and Shigeki Aoki. 2016. “Prospective estimation of mean axon diameter and extra-axonal space of the posterior limb of the internal capsule in patients with idiopathic normal pressure hydrocephalus before and after a lumboperitoneal shunt by using q-space diffusion MRI.” European Radiology 26 (9): 2992–2998. https://doi.org/10.1007/s00330-015-4162-9.

Jensen, Jens H, Joseph A Helpern, Anita Ramani, Hanzhang Lu, and Kyle Kaczynski. 2005. “Diffusional kurtosis imaging: the quantification of non–gaussian water diffusion by means of magnetic resonance imaging.” Magnetic Resonance in Medicine: An Official Journal of the International Society for Magnetic Resonance in Medicine 53 (6): 1432–1440.

Jones, Robert, Giorgia Grisot, Jean Augustinack, Caroline Magnain, David A. Boas, Bruce Fischl, Hui Wang, and Anastasia Yendiki. 2020. “Insight into the fundamental trade-offs of diffusion MRI from polarization-sensitive optical coherence tomography in ex vivo human brain.” NeuroImage 214: 116704. https://doi.org/https://doi.org/10.1016/j.neuroimage.2020.116704. http://www.sciencedirect.com/science/article/pii/S1053811920301919.

Jones, Robert, Qiyuan Tian, Chiara Maffei, Jean Augustinack, Aapo Nummenmaa, Susie Huang, and Anastasia Yendiki. 2021. “Generalized anisotropy profiles distinguish cortical and subcortical structures in ex vivo diffusion MRI.” Proc. Intl. Soc. Mag. Res. Med.

Keil, Boris, James N. Blau, Stephan Biber, Philipp Hoecht, Veneta Tountcheva, Kawin Setsompop, Christina Triantafyllou, and Lawrence L. Wald. 2013. “A 64-channel 3T array coil for accelerated brain MRI.” Magnetic Resonance in Medicine 70 (1): 248–258. https://doi.org/10.1002/mrm.24427. https://onlinelibrary.wiley.com/doi/abs/10.1002/mrm.24427.

Khan, Ahmad Raza, Anda Cornea, Lindsey A Leigland, Steven G Kohama, Sune Nørhøj Jespersen, and Christopher D Kroenke. 2015. “3D structure tensor analysis of light microscopy data for validating diffusion MRI.” Neuroimage 111: 192–203.

Lacerda, Luis M, Jonathan I Sperl, Marion I Menzel, Tim Sprenger, Gareth J Barker, and Flavio Dell'Acqua. 2016. “Diffusion in realistic biophysical systems can lead to aliasing effects in diffusion spectrum imaging.” Magnetic resonance in medicine 76 (6): 1837–1847.

Le Bihan, Denis, Eric Breton, Denis Lallemand, Philippe Grenier, Emmanuel Cabanis, and Maurice Laval-Jeantet. 1986. “MR imaging of intravoxel incoherent motions: application to diffusion and perfusion in neurologic disorders.” Radiology 161 (2): 401–407.

Leergaard, Trygve B, Nathan S White, Alex De Crespigny, Ingeborg Bolstad, Helen D'Arceuil, Jan G Bjaalie, and Anders M Dale. 2010. “Quantitative histological validation of diffusion MRI fiber orientation distributions in the rat brain.” PloS one 5 (1): e8595.

Lefebvre, Joël, Patrick Delafontaine-Martel, Philippe Lemieux, Maxime Descoteaux, Laurent Petit, and Frédéric Lesage. 2021. “Localization and imaging of white matter fiber crossings in whole mouse brains using diffusion MRI and serial blockface OCT.” Optical Techniques in Neurosurgery, Neurophotonics, and Optogenetics.

Liewald, Daniel, Robert Miller, Nikos Logothetis, Hans-Joachim Wagner, and Almut Schüz. 2014. “Distribution of axon diameters in cortical white matter: an electron-microscopic study on three human brains and a macaque.” Biological cybernetics 108 (5): 541–557. https://doi.org/10.1007/s00422-014-0626-2. https://www.ncbi.nlm.nih.gov/pubmed/25142940. https://www.ncbi.nlm.nih.gov/pmc/articles/PMC4228120/.

Liu, Chao J, Adam J Black, Hui Wang, and Taner Akkin. 2016. “Quantifying three-dimensional optic axis using polarization-sensitive optical coherence tomography.” Journal of biomedical optics 21 (7): 070501.

Lustig, Michael, David L Donoho, Juan M Santos, and John M Pauly. 2008. “Compressed sensing MRI.” IEEE signal processing magazine 25 (2): 72–82.

Lustig, Michael, David Donoho, and John M Pauly. 2007. “Sparse MRI: The application of compressed sensing for rapid MR imaging.” Magnetic Resonance in Medicine: An Official Journal of the International Society for Magnetic Resonance in Medicine 58 (6): 1182–1195.

Maffei, C., G. Girard, K. G. Schilling, N. Adluru, D. B. Aydogan, A. Hamamci, F.-C. Yeh, M. Mancini, Y. Wu, A. Sarica, A. Teillac, S. H. Baete, D. Karimi, Y.-C. Lin, F. Boada, N. Richard, B. Hiba, A. Quattrone, Y. Hong, D. Shen, P.-T. Yap, T. Boshkovski, J. S. W. Campbell, N. Stikov, G. B. Pike, B. B. Bendlin, A. L. Alexander, V. Prabhakaran, A. Anderson, B. A. Landman, E. J. Z. Canales-Rodríguez, M. Barakovic, J. Rafael-Patino, T. Yu, G. Rensonnet, S. Schiavi, A. Daducci, M. Pizzolato, E. Fischi-Gomez, J.-P. Thiran, G. Dai, G. Grisot, N. Lazovski, A. Puente, M. Rowe, I. Sanchez, V. Prchkovska, R. Jones, J. Lehman, S. Haber, and A. Yendiki. 2020. “The IronTract challenge: Validation and optimal tractography methods for the HCP diffusion acquisition scheme.” ISMRM (oral presentation).

Maffei, C., G. Girard, K. G. Schilling, N. Adluru, D. B. Aydogan, A. Hamamci, F.-C. Yeh, M. Mancini, Y. Wu, A. Sarica, A. Teillac, S. H. Baete, D. Karimi, Y.-C. Lin, F. Boada, N. Richard, B. Hiba, A. Quattrone, Y. Hong, D. Shen, P.-T. Yap, T. Boshkovski, J. S. W. Campbell, N. Stikov, G. B. Pike, B. B. Bendlin, A. L. Alexander, V. Prabhakaran, A. Anderson, B. A. Landman, E. J. Z. Canales-Rodríguez, M. Barakovic, J. Rafael-Patino, T. Yu, G. Rensonnet, S. Schiavi, A. Daducci, M. Pizzolato, E. Fischi-Gomez, J.-P. Thiran, G. Dai, G. Grisot, N. Lazovski, A. Puente, M. Rowe, I. Sanchez, V. Prchkovska, R. Jones, J. Lehman, S. Haber, and A. Yendiki. 2021. “New insights from the IronTract challenge: Simple post-processing enhances the accuracy of diffusion tractography.” Proc Int Soc Magn Reson Med (ISMRM).

Majka, Piotr, and Daniel K Wójcik. 2016. “Possum—a framework for three-dimensional reconstruction of brain images from serial sections.” Neuroinformatics 14 (3): 265–278.

Menzel, Marion I, Ek T Tan, Kedar Khare, Jonathan I Sperl, Kevin F King, Xiaodong Tao, Christopher J Hardy, and Luca Marinelli. 2011. “Accelerated diffusion spectrum imaging in the human brain using compressed sensing.” Magnetic Resonance in Medicine 66 (5): 1226–1233.

Mollink, Jeroen, Michiel Kleinnijenhuis, Anne-Marie van Cappellen van Walsum, Stamatios N Sotiropoulos, Michiel Cottaar, Christopher Mirfin, Mattias P Heinrich, Mark Jenkinson, Menuka Pallebage-Gamarallage, and Olaf Ansorge. 2017. “Evaluating fibre orientation dispersion in white matter: comparison of diffusion MRI, histology and polarized light imaging.” Neuroimage 157: 561–574.

Paquette, Michael, Guillaume Gilbert, and Maxime Descoteaux. 2016. “Optimal DSI reconstruction parameter recommendations: better ODFs and better connectivity.” NeuroImage 142: 1–13.

Paquette, Michael, Sylvain Merlet, Guillaume Gilbert, Rachid Deriche, and Maxime Descoteaux. 2015. “Comparison of sampling strategies and sparsifying transforms to improve compressed sensing diffusion spectrum imaging.” Magnetic resonance in medicine 73 (1): 401–416.

Preibisch, Stephan, Stephan Saalfeld, and Pavel Tomancak. 2009. “Globally optimal stitching of tiled 3D microscopic image acquisitions.” Bioinformatics 25 (11): 1463–1465.

Reese, Timothy G, Thomas Benner, Ruopeng Wang, David A Feinberg, and Van J Wedeen. 2009. “Halving imaging time of whole brain diffusion spectrum imaging and diffusion tractography using simultaneous image refocusing in EPI.” Journal of Magnetic Resonance Imaging: An Official Journal of the International Society for Magnetic Resonance in Medicine 29 (3): 517–522.

Reuter, Martin, H Diana Rosas, and Bruce Fischl. 2010. “Highly accurate inverse consistent registration: a robust approach.” Neuroimage 53 (4): 1181–1196.

Ruiz-Lopera, Sebastián, René Restrepo, Taylor M Cannon, Martin Villiger, Brett E Bouma, and Néstor Uribe-Patarroyo. 2021. “Computational refocusing in polarization-sensitive optical coherence tomography with phase unstable systems.” Optical Coherence Tomography and Coherence Domain Optical Methods in Biomedicine XXV.

Schilling, Kurt G, Vaibhav Janve, Yurui Gao, Iwona Stepniewska, Bennett A Landman, and Adam W Anderson. 2018. “Histological validation of diffusion MRI fiber orientation distributions and dispersion.” Neuroimage 165: 200–221.

Schilling, Kurt, Yurui Gao, Vaibhav Janve, Iwona Stepniewska, Bennett A Landman, and Adam W Anderson. 2017. “Can increased spatial resolution solve the crossing fiber problem for diffusion MRI.” NMR in Biomedicine 30 (12): e3787.

Schilling, Kurt, Vaibhav Janve, Yurui Gao, Iwona Stepniewska, Bennett A Landman, and Adam W Anderson. 2016. “Comparison of 3D orientation distribution functions measured with confocal microscopy and diffusion MRI.” Neuroimage 129: 185–197.

Schindelin, Johannes, Ignacio Arganda-Carreras, Erwin Frise, Verena Kaynig, Mark Longair, Tobias Pietzsch, Stephan Preibisch, Curtis Rueden, Stephan Saalfeld, and Benjamin Schmid. 2012. “Fiji: an open-source platform for biological-image analysis.” Nature methods 9 (7): 676.

Seehaus, Arne, Alard Roebroeck, Matteo Bastiani, Lúcia Fonseca, Hansjürgen Bratzke, Nicolás Lori, Anna Vilanova, Rainer Goebel, and Ralf Galuske. 2015. “Histological validation of high-resolution DTI in human post mortem tissue.” Frontiers in Neuroanatomy 9: 98.

Setsompop, K., R. Kimmlingen, E. Eberlein, T. Witzel, J. Cohen-Adad, J. A. McNab, B. Keil, M. D. Tisdall, P. Hoecht, P. Dietz, S. F. Cauley, V. Tountcheva, V. Matschl, V. H. Lenz, K. Heberlein, A. Potthast, H. Thein, J. Van Horn, A. Toga, F. Schmitt, D. Lehne, B. R. Rosen, V. Wedeen, and L. L. Wald. 2013. “Pushing the limits of in vivo diffusion MRI for the Human Connectome Project.” NeuroImage 80: 220–233. https://doi.org/10.1016/j.neuroimage.2013.05.078. https://www.ncbi.nlm.nih.gov/pubmed/23707579. https://www.ncbi.nlm.nih.gov/pmc/articles/PMC3725309/. https://www.ncbi.nlm.nih.gov/pmc/articles/PMC3725309/pdf/nihms484864.pdf.

Setsompop, Kawin, Julien Cohen-Adad, Borian A Gagoski, Tommi Raij, Anastasia Yendiki, Boris Keil, Van J Wedeen, and Lawrence L Wald. 2012. “Improving diffusion MRI using simultaneous multi-slice echo planar imaging.” Neuroimage 63 (1): 569–580.

Setsompop, Kawin, Qiuyun Fan, Jason Stockmann, Berkin Bilgic, Susie Huang, Stephen F Cauley, Aapo Nummenmaa, Fuyixue Wang, Yogesh Rathi, and Thomas Witzel. 2018. “High– resolution in vivo diffusion imaging of the human brain with generalized slice dithered enhanced resolution: Simultaneous multislice (g S lider–SMS).” Magnetic resonance in medicine 79 (1): 141–151.

Setsompop, Kawin, Borjan A Gagoski, Jonathan R Polimeni, Thomas Witzel, Van J Wedeen, and Lawrence L Wald. 2012. “Blipped–controlled aliasing in parallel imaging for simultaneous multislice echo planar imaging with reduced g–factor penalty.” Magnetic resonance in medicine 67 (5): 1210–1224.

Stejskal, Edward O, and John E Tanner. 1965. “Spin diffusion measurements: spin echoes in the presence of a time–dependent field gradient.” The journal of chemical physics 42 (1): 288–292.

Tian, Qiyuan, Ariel Rokem, Rebecca D Folkerth, Aapo Nummenmaa, Qiuyun Fan, Brian L Edlow, and Jennifer A McNab. 2016. “Q–space truncation and sampling in diffusion spectrum imaging.” Magnetic resonance in medicine 76 (6): 1750–1763.

Tobisch, Alexandra, Rüdiger Stirnberg, Robbert Leonard Harms, Thomas Schultz, Alard Roebroeck, Monique Breteler, and Tony Stöcker. 2018. “Compressed sensing diffusion spectrum imaging for accelerated diffusion microstructure MRI in long-term population imaging.” Frontiers in neuroscience 12: 650.

Tuch, David S. 2004. “Q–ball imaging.” Magnetic Resonance in Medicine: An Official Journal of the International Society for Magnetic Resonance in Medicine 52 (6): 1358–1372.

Ugryumova, Nadezhda, Sergei V Gangnus, and Stephen J Matcher. 2006. “Three-dimensional optic axis determination using variable-incidence-angle polarization-optical coherence tomography.” Optics letters 31 (15): 2305–2307.

Ugryumova, Nadya, James Jacobs, Marco Bonesi, and Stephen J Matcher. 2009. “Novel optical imaging technique to determine the 3-D orientation of collagen fibers in cartilage: variable-incidence angle polarization-sensitive optical coherence tomography.” Osteoarthritis and cartilage 17 (1): 33–42.

Villiger, Martin, Boy Braaf, Norman Lippok, Kenichiro Otsuka, Seemantini K Nadkarni, and Brett E Bouma. 2018. “Optic axis mapping with catheter-based polarization-sensitive optical coherence tomography.” Optica 5 (10): 1329–1337.

Wang, Hui, Taner Akkin, Caroline Magnain, Ruopeng Wang, Jay Dubb, William J Kostis, Mohammad A Yaseen, Avilash Cramer, Sava Sakadžić, and David Boas. 2016. “Polarization sensitive optical coherence microscopy for brain imaging.” Optics letters 41 (10): 2213–2216.

Wang, Hui, Adam J Black, Junfeng Zhu, Tyler W Stigen, Muhammad K Al-Qaisi, Theoden I Netoff, Aviva Abosch, and Taner Akkin. 2011. “Reconstructing micrometer-scale fiber pathways in the brain: multi-contrast optical coherence tomography based tractography.” Neuroimage 58 (4): 984–992.

Wang, Hui, Christophe Lenglet, and Taner Akkin. 2015. “Structure tensor analysis of serial optical coherence scanner images for mapping fiber orientations and tractography in the brain.” Journal of biomedical optics 20 (3): 036003.

Wang, Hui Caroline Magnain, Ruopeng Wang, Jay Dubb, Ani Varjabedian, Lee S Tirrell, Allison Stevens, Jean C Augustinack, Ender Konukoglu, and Iman Aganj. 2018. “as-PSOCT: Volumetric microscopic imaging of human brain architecture and connectivity.” Neuroimage 165: 56–68.

Wang, Hui, Junfeng Zhu, Martin Reuter, Louis N Vinke, Anastasia Yendiki, David A Boas, Bruce Fischl, and Taner Akkin. 2014. “Cross-validation of serial optical coherence scanning and diffusion tensor imaging: a study on neural fiber maps in human medulla oblongata.” Neuroimage 100: 395–404.

Wedeen, Van J, Patric Hagmann, Wen-Yih Isaac Tseng, Timothy G Reese, and Robert M Weisskoff. 2005. “Mapping complex tissue architecture with diffusion spectrum magnetic resonance imaging.” Magnetic resonance in medicine 54 (6): 1377–1386.

Wedeen, Van J, RP Wang, Jeremy D Schmahmann, Thomas Benner, Wen-Yih Isaac Tseng, Guangping Dai, DN Pandya, Patric Hagmann, Helen D'Arceuil, and Alex J de Crespigny. 2008. “Diffusion spectrum magnetic resonance imaging (DSI) tractography of crossing fibers.” Neuroimage 41 (4): 1267–1277.

Yao, Gang, and Dongsheng Duan. 2020. “High-resolution 3D tractography of fibrous tissue based on polarization-sensitive optical coherence tomography.” Experimental Biology and Medicine 245 (4): 273–281.

Yeh, Fang-Cheng, Van Jay Wedeen, and Wen-Yih Isaac Tseng. 2010. “Generalized ${q} $-Sampling Imaging.” IEEE transactions on medical imaging 29 (9): 1626–1635.

Yendiki, A., R. Jones, A. Dalca, H. Wang, and B. Fischl. 2020. “Towards taking the guesswork (and the errors) out of diffusion tractography.” Proc Int Soc Magn Reson Med (ISMRM).

Yendiki, Anastasia, Manisha Aggarwal, Markus Axer, Amy F. D. Howard, Anne-Marie van Cappellen van Walsum, and Suzanne N. Haber. 2021. “Post mortem mapping of connectional anatomy for the validation of diffusion MRI.” bioRxiv: 2021.04.16.440223. https://doi.org/10.1101/2021.04.16.440223. https://www.biorxiv.org/content/biorxiv/early/2021/04/19/2021.04.16.440223.full.pdf.

Zhang, Hui, Torben Schneider, Claudia A Wheeler-Kingshott, and Daniel C Alexander. 2012. “NODDI: practical in vivo neurite orientation dispersion and density imaging of the human brain.” Neuroimage 61 (4): 1000–1016.

